# The African Swine Fever Virus gene MGF_360-4L inhibits interferon signaling by recruiting mitochondrial selective autophagy receptor SQSTM1 degrading MDA5 antagonizing innate immune responses

**DOI:** 10.1101/2024.09.09.612163

**Authors:** Hualin Sun, Jifei Yang, Zhonghui Zhang, Mengli Wu, Zhancheng Tian, Ying Liu, Xiaoqiang Zhang, Jianhao Zhong, Songlin Yang, Yikang Chen, Jianxun Luo, Guiquan Guan, Hong Yin, Qingli Niu

## Abstract

Multigene family (MGF) 360 genes, which are African swine fever virus (ASFV) virulence genes, primarily target key host immune molecules to suppress host interferon (IFN) production and interferon-stimulated gene (ISG) transcription, impairing host innate immune responses for efficient viral replication. However, the interactions between MGF 360 virulence genes and host molecules, as well as the mechanisms through which MGF 360 genes regulate host immune responses and interferon signaling, require further elucidation. In this study, we discovered that ASFV MGF_360-4L interacts with MDA5 and recruits the mitochondrial selective autophagy receptor SQSTM1 to degrade MDA5, thus impairing interferon signaling and compromising host innate immune responses. Furthermore, MGF_360-4L inhibits the interaction between MDA5 and MAVS, blocking ISG15-mediated ISGylation of MDA5. MGF_360-4L deficiency significantly attenuated virus-induced mitochondrial autophagy *in vitro*. Additionally, OAS1 ubiquitinates MGF_360-4L at residues K290, K295 and K327. Finally, a recombinant ASFV lacking the MGF_360-4L gene (ASFV-ΔMGF_360-4L) was generated using ASFV-CN/SC/2019 as the backbone, which demonstrated that the replication kinetics of ASFV-ΔMGF_360-4L in PAM cells were like those of the highly virulent parental ASFV-WT *in vitro*. Domestic pigs infected with ASFV-ΔMGF_360-4L exhibited milder symptoms than those infected with parental ASFV-WT, and ASFV-ΔMGF_360-4L-infected pigs presented with enhanced host innate antiviral immune response, confirming that the deletion of the MGF_360-4L gene from the ASFV genome highly attenuated virulence in pigs and provided effective protection against parental ASFV challenge. In conclusion, we identified a novel ASFV virulence gene, MGF_360-4L, further elucidating ASFV infection mechanisms and providing a new candidate for vaccine development.

**IMPORTANCE:** African swine fever virus (ASFV) infection causes acute death in pigs, and there is currently no effective vaccine available for prevention. Multigene family (MGF) virulence genes have been shown to be crucial for ASFV ability to evade host innate immune responses. However, the functions of most MGF genes remain unknown, which poses significant challenges for the development of ASFV vaccines and antiviral drugs. In this study, we identified a virulence gene of ASFV, MGF_360-4L, that targets and recruits the selective autophagy receptor p62 to mediate the degradation of the dsRNA sensor MDA5, thereby blocking interferon signaling. Additionally, it inhibits the ISG15-mediated ISGylation activation of MDA5. ASFV lacking MGF_360-4L showed reduced virulence and provided protection in pigs. Our data identify a novel virulence gene and provide new insights for ASFV vaccine development.

## Introduction

African swine fever (ASF) is a highly contagious and fatal infectious disease in pigs caused by African swine fever virus (ASFV), which is the sole member of the *Asfarviridae* family and a large icosahedral DNA arbovirus (1). It spreads not only through direct contact but also via soft tick bites (2). The lethality of ASF is closely linked to viral pathogenicity, with high morbidity and mortality rates reaching 100%, causing serious economic consequences for the swine industry (3). ASF was first identified in Kenya in 1921 and subsequently spread rapidly from Africa to the Caucasus (4-6). In 2007, it reached the Republic of Georgia and spread to most countries in the eastern part of the European Union in 2014 (7). In 2018, it was first reported in China (8), followed by Vietnam (9) and South Korea (10). Owing to the large ASFV genome and complex pathogenesis, different vaccine strategies for ASF have been evaluated in recent decades. To date, there is still a lack of safe and effective vaccines, and ASF continues to pose a global threat to the swine industry, severely impacting its security (3,11).

ASFV has a genome ranging from 170 to 193 kb, encoding 150-200 proteins, many of whose functions remain largely unknown (12). During viral infection, pathogen-associated molecular patterns (PAMPs) are recognized by cytosolic pattern recognition receptors (PRRs), activating innate immune responses that induce the production of inflammatory factors and interferons (IFNs) to combat viral infections (13). IFNs are primarily induced by RIG-I/MDA5, cGAS-STING and Toll-like receptors, followed by the transcription of interferon-stimulated genes (ISGs) to further suppress viral infections (14-17). The specific PRRs involved in ASFV infection are largely unclear, and some studies suggest that the ASFV genome is rich in AT sequences recognized by RNA Pol-III, leading to RIG-I-mediated innate immune responses. However, another study indicated that the ASFV virulence gene I267L impairs RIG-I activation and stability, thus inhibiting IFN signaling (18).

MDA5 and RIG-I are conserved cytosolic PRRs that share N-terminal tandem caspase activation and recruitment domains (CARDs). These domains interact with the mitochondrial antiviral signaling protein MAVS (also known as IPS-1, VISA, and Cardif) (19), which is located on the outer mitochondrial membrane via its C-terminal transmembrane domain and interacts with the CARD domains of RIG-I/MDA5 to induce interferon signaling (20). TBK1/IKKε are subsequently activated and lead to the phosphorylation and nuclear translocation of IRF3 and IRF7, stimulating interferon secretion (21). Several studies have reported that viral infection induces mitochondrial autophagy to degrade MAVS, suppressing innate immune responses and promoting viral replication (22-25). The activation of MDA5 can also depend on ISG15-mediated ISGylation of CARD domains, stimulating innate immune responses, which are specific to MDA5 activation rather than RIG-I (26). However, the function of ASFV genes that target MDA5 and the related mechanisms by which ASFV affects MDA5 remain unclear.

Autophagy plays a crucial role in cellular processes and maintaining homeostasis and involves the selective degradation of damaged organelles and misfolded proteins and the capture of pathogens within autophagolysosomes (27). Autophagy is selective, targeting specific substrates via autophagy receptors such as SQSTM1/p62, TOLLIP, NDP52, NBR1, OPTN, and CCDC50 (28,29). During ASFV infection, multiple gene family (MGF) proteins promote autophagy to counteract innate immune responses. For example, the MGF-300-2R protein recruits TOLLIP to promote IKKα and IKKβ autophagic degradation, impairing innate immune responses (30). The pA137R protein interacts with and targets TBK1 for degradation via the autophagolysosomal pathway, inhibiting IRF3 nuclear translocation and affecting cGAS-STING signaling activation (31). The MGF-505-7R protein interacts with STING and increases the expression of the autophagy-related protein ULK1 to degrade STING, suppressing cGAS-STING signaling activation (32). Moreover, L83L interacts with cGAS and STING, recruiting TOLLIP to facilitate the autophagic degradation of STING, thereby blocking downstream signaling and reducing IFN-I production (33). Additionally, the p17 protein induces mitochondrial autophagy through interaction with SQSTM1 (P62) and TOMM70, modulating innate immunity (34). MGF proteins can also inhibit host innate immunity by promoting ubiquitination. For example, the MGF-360-9L protein, a virulence factor, degrades STAT1 and STAT2 via ubiquitination, suppressing IFN-β secretion (35). MGF-360-10L recruits the E3 ubiquitin ligase HERC5 to mediate K48-linked ubiquitination and degradation of JAK1 (36). However, further research is needed to fully understand the roles of MGF proteins in inducing autophagy and regulating host immune responses and signaling pathways. Additionally, further investigations are needed to elucidate the detailed mechanisms by which MGF proteins regulate autophagy to influence host immune responses and signaling pathways during ASFV infection.

In previous studies, we demonstrated that OAS1 serves as a critical antiviral factor by promoting TRIM21-mediated ubiquitination to increase the proteasomal degradation of P72, thereby inhibiting ASFV replication and stimulating innate immune activation (37). Furthermore, we found that OAS1 interacts not only with P72 but also with several MGF proteins, suggesting potential underlying mechanisms for OAS1. However, these mechanisms require further investigation. In this study, we revealed that MGF_360-4L interacts with MDA5, inducing mitochondrial autophagy mediated by the p62-selective autophagy receptor, leading to MDA5 degradation, suppression of interferon signaling, and inhibition of MDA5 ISGylation activation. Subsequent research revealed that OAS1, as a viral restriction factor, facilitates the ubiquitination and degradation of MGF_360-4L at residues K290, K295, and K327, thereby attenuating the inhibitory effect of MGF_360-4L on MDA5. Our findings establish MGF_360-4L as a crucial virulence factor that exploits autophagy to modulate host immune responses and promote ASFV replication. Furthermore, MGF_360-4L gene deficiency attenuated ASFV pathogenicity in pigs. These discoveries are pivotal for understanding how ASFV evades host immune responses and for developing potential therapeutic strategies.

## Results

### The ASFV MGF_360-4L protein inhibits RIG-I/MDA5-mediated activation of the IFN signaling pathway

Previous studies have indicated that ASFV infection induces autophagy while suppressing interferon production (30-33). However, there are no reports on the regulatory effects of MGF_360-4L on the host. In our study, we initially predicted that OAS1 might interact with MGF_360-4L based on Co-IP/LC-MS/MS data (Fig. S1). Given that OAS1 is an interferon-stimulated gene (ISG), we speculate that MGF_360-4L may influence the interferon signaling pathway. To further elucidate the regulatory effect of ASFV MGF_360-4L on the activation of IFN promoters, ISRE promoters or the NF-κB promoter, as well as the changes in the expression of molecules related to the interferon signaling pathway specifically targeted by MGF_360-4L, HEK293T cells were cotransfected with a plasmid expressing MGF_360-4L, along with RIG-I, MDA5, MAVS/VISA, TRAF3, IKKε, cGAS, STING/MITA, TBK1, IRF3, or IRF7 luciferase reporters, and a reporter assay was conducted. Plasmids expressing the dsRNA mimic poly(I:C) and the dsDNA mimic poly(dA:dT) were used as controls. As shown in Fig.1A-C, compared with the dsRNA mimic poly(I:C) and the dsDNA mimic poly(dA:dT), MGF_360-4L suppressed the activation of the IFN-β promoter and the ISRE promoter but not the NF-κB promoter. Further analysis revealed that MGF_360-4L inhibited the activation of the IFN-β promoter stimulated by poly(I:C) and poly(dA:dT) and suppressed the activation of MAVS/VISA, TRAF3, IKKε, TBK1, IRF3, and IRF7, which are downstream molecules of MDA5 (Fig.1D). However, MGF_360-4L did not affect the activation of IFN-β triggered by cGAS or STING/MITA (Fig.1E). RT‒qPCR analysis revealed that MGF_360-4L significantly inhibited the transcript levels of IFIT1, IFIT2, STAT1, STAT2, MX1, and OAS1, which are regulated by IFN-β, but not OASL. Interestingly, MGF_360-4L also increased the transcript level of ISG15, and this effect was further enhanced in the presence of IFN-β (Fig.1F). Collectively, these results suggest that MGF_360-4L negatively regulates the activation of IFN-β mediated by RIG-I/MDA5.

**Fig 1.**
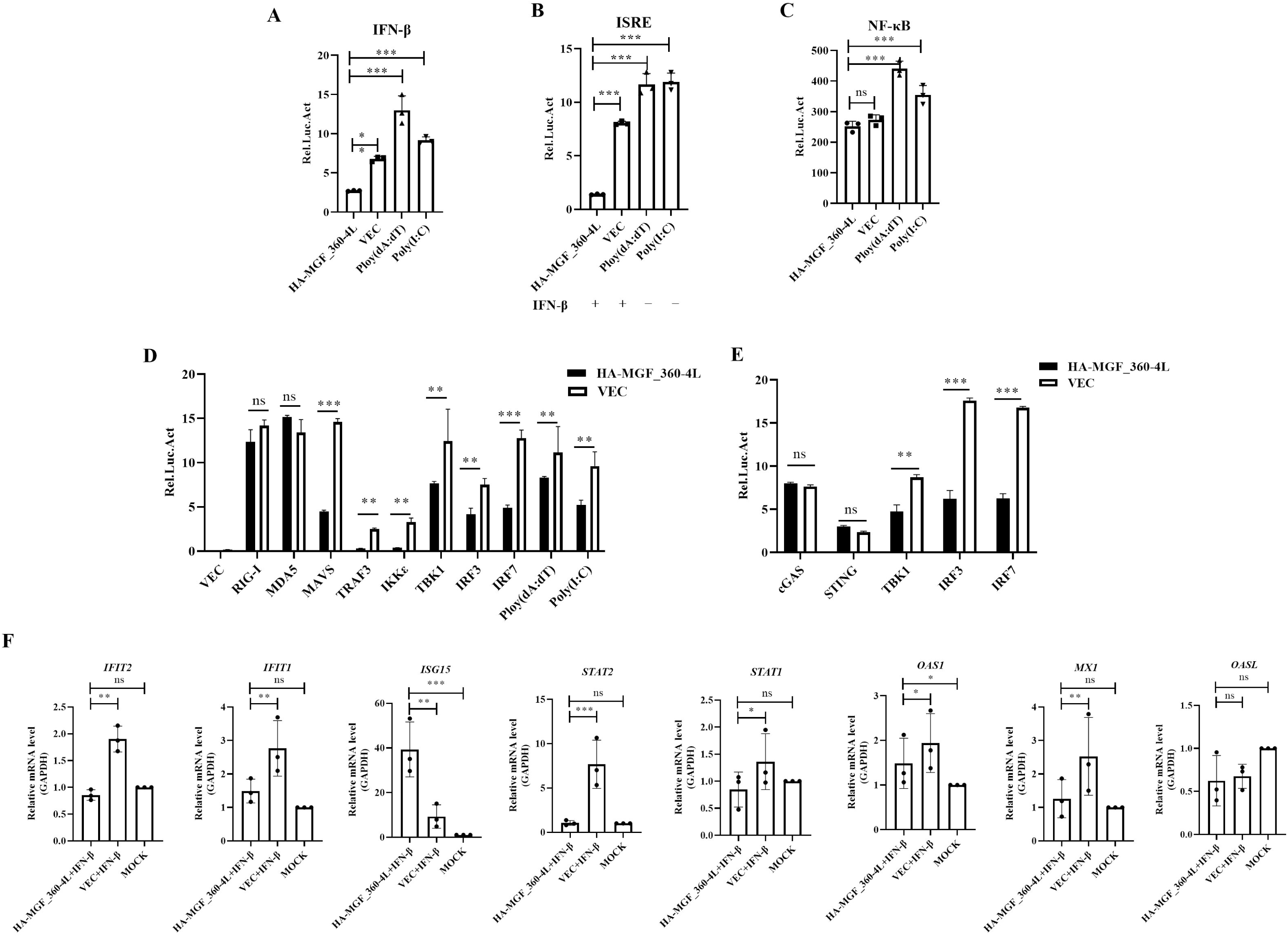
MGF_360-4L inhibits INF-β and ISRE promoter activity while activating ISG transcription. (A-C) HEK293T cells were transfected with plasmids carrying the IFN-β, ISRE, or NF-κB promoters, along with the TK plasmid and HA-MGF_360-4L plasmid or empty vector. Cells were stimulated with poly(dA:dT) or poly(I:C) as controls, and the fluorescence intensities were measured to assess promoter activation. (D-E) HEK293T cells were transfected with plasmids carrying the IFN-β promoter and TK plasmid, along with HA-tagged plasmids (RIG-I, MDA5, MAVS/VISA, TRAF3, IKKε, cGAS, STING/MITA, TBK1, IRF3, and IRF7) and HA-MGF_360-4L plasmid or empty vector. Cells were stimulated with poly(dA:dT) or poly(I:C) as controls, and the fluorescence intensities were measured to assess promoter activation. (F) ISG transcript levels were measured in HEK293T cells transfected with HA-MGF_360-4L or empty vector and stimulated with or without IFN-β.

### The MGF_360-4L protein interacts with MDA5

The results presented above indicate that MGF_360-4L functions upstream of RIG-I/MDA5 in the IFN signaling pathway. To investigate how MGF_360-4L regulates RIG-I/MDA5-mediated IFN signaling, coimmunoprecipitation (Co-IP) of MGF_360-4L with various mediators, including RIG-I, MDA5, MAVS/VISA, TRAF3, IKKε, TBK1, IRF3, and IRF7 of the IFN signaling pathway, was performed. Co-IP results indicated that Flag-MGF_360-4L specifically interacts with HA-MDA5 but not with other molecules (Fig.2A-B). Confocal microscopy revealed that MGF_360-4L and MDA5 colocalize in the cytoplasm of cells (Fig.2C). Additionally, GST pull-down assays further demonstrated that His-MDA5 was pulled down by GST-MGF_360-4L (Fig.2D). To further determine the structural domains involved in the interaction between MGF_360-4L and MDA5, we constructed Flag-tagged MDA5 plasmids for the following domains: CARD_MDA5_R1 (9--100 aa), CARD_MDA5_R2 (112--202 aa), Type III restriction enzyme (257--443 aa), MDA5 insert domain (500--640 aa), helicase_C domain (700--839 aa), and MDA5-C domain (905--1021 aa) (38). Simultaneously, using the Protein Crystal Database to evaluate the protein structure, we constructed three fragments of MGF_360-4L, which were dimeric DARP in complex with EpoR (34--92 aa), the RNASEL complex with sunitinib (66--118 aa) and Alpha-lactotoxin_LT1a (191--337 aa), via SWISS-MODEL online simulation (https://swissmodel.ExPASy.org/interactive) (Fig.2E). Co-IP results revealed that the Alpha-lactotoxin_LT1a-like domain of MGF_360-4L specifically interacts with MDA5 and that the Helicase_C domain of MDA5 specifically interacts with MGF_360-4L (Fig.2F-G). Confocal microscopy also revealed that when either the Alpha-lactotoxin_LT1a-like domain of MGF_360-4L or the SF2_C dicer domain of MDA5 was deleted, colocalization between wild-type MDA5 and MGF_360-4L could not be observed in cells (Fig.2H). These results indicate that MGF_360-4L specifically interacts with MDA5 to counteract the activation of the IFN signaling pathway.

**Fig 2.**
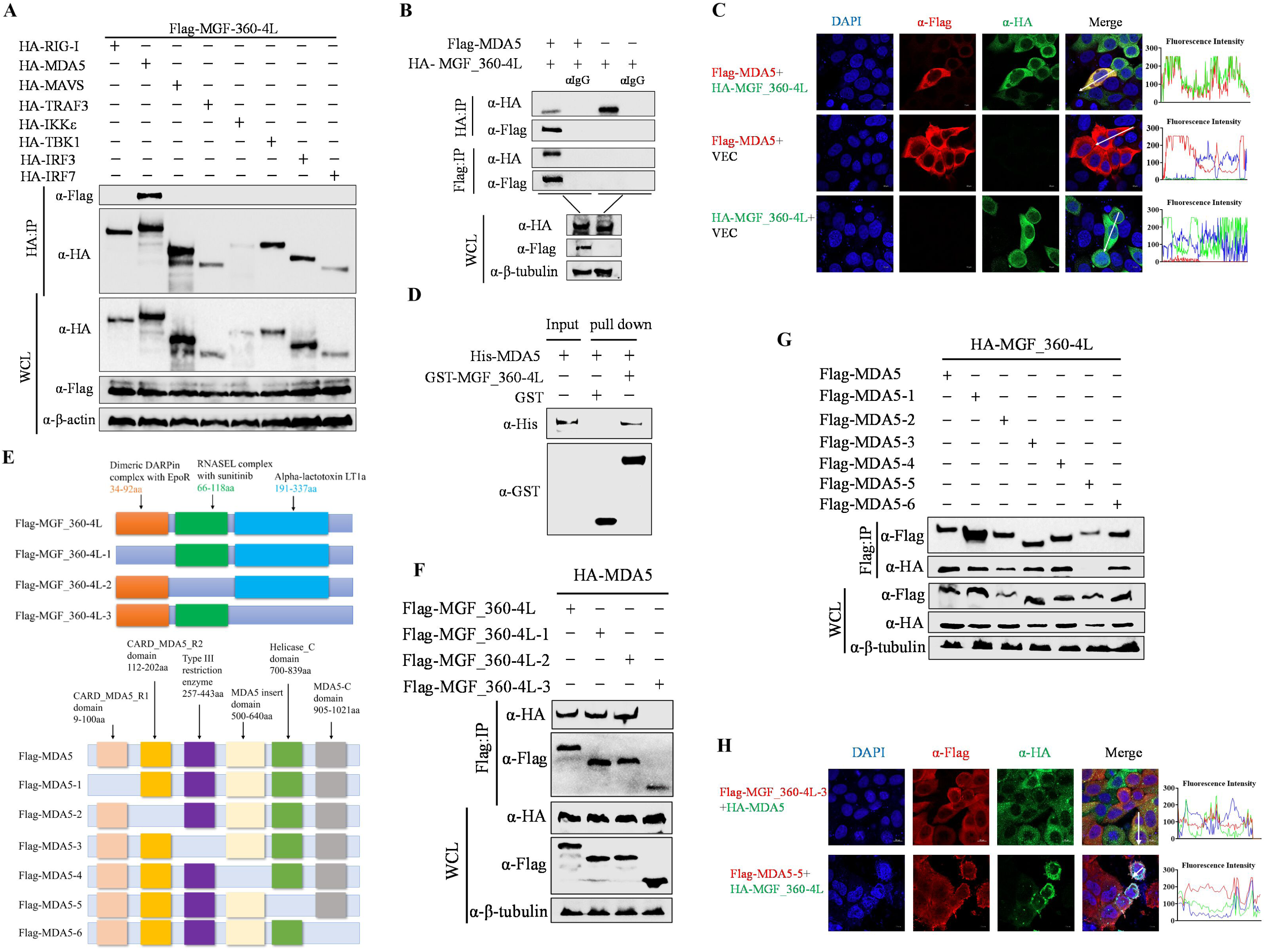
The MGF_360-4L protein interacts with MDA5. (A-B) HEK293T cells were cotransfected with plasmids expressing Flag-MGF_360-4L and HA-RIG-I, MDA5, MAVS/VISA, TRAF3, IKKε, TBK1, IRF3 or IRF7 as indicated. Co-IP analysis was performed to detect the interaction between MGF_360-4L and RIG-I, MDA5, MAVS/VISA, TRAF3, IKKε, TBK1, IRF3 or IRF7 after 24 h. (C) HeLa cells were transfected with pHA-MGF_360-4L and pFlag-MDA5 or pHA-MGF_360-4L or pFlag-MDA5 with empty vector VEC. The localization of MGF_360-4L and MDA5 was detected via immunofluorescence microscopy. Scale bars, 10 μm. (D) GST-tagged MGF_360-4L protein (10 μg) or GST antigen was added to GST-Sepharose 4B beads, followed by binding with His-tagged MDA5 protein (10 μg). The samples were subjected to Western blotting analysis to detect direct interactions between MGF_360-4L-GST and MDA5-His. (E) Schematic diagram of the construction of MGF_360-4L and MDA5 fragments. (F-G) HEK293T cells were cotransfected with Flag-MGF_360-4L (1-3) truncations and pHA-MDA5 (F) or Flag-MDA5 (1-6) truncations and HA-MGF_360-4L (G). Co-IP analysis was performed to detect the interaction between MGF_360-4L and MDA5. (H) HeLa cells were transfected with pFlag-MGF_360-4L-3 and pHA-MDA5 or pFlag-MDA5-5 and pHA-MGF_360-4L. The localizations of MGF_360-4L-3 and MDA5 or MDA5-5 and MGF_360-4L were detected via immunofluorescence microscopy. Scale bars, 10 μm. Fluorescence intensity profiles were measured along the drawn measurement line.

### MGF_360-4L inhibits the MDA5-mediated interferon signaling pathway and IRF3 nuclear ***translocation***

Next, we further investigated how MGF_360-4L suppresses the interferon signaling pathway, thereby limiting the innate immune response. First, we transfected HA-MGF_360-4L or HA-Vec plasmids into HET293T cells and stimulated them with vesicular stomatitis virus (VSV) at an MOI of 1 for 0, 3, 6, or 9 hours postinfection (hpi). The Western blot results indicated that MGF_360-4L specifically reduced the MDA5 protein level and inhibited the phosphorylation of IKKε, TBK1, IRF3 and IRF7 (Fig.3A). Additionally, nucleocytoplasmic separation assays further confirmed that MGF_360-4L obstructed IRF3 nuclear translocation; interestingly, MGF_360-4L was also found to partially enter the nucleus (Fig.3B). Confocal microscopy results confirmed that MGF_360-4L affects the nuclear translocation of IRF3 (Fig.3C). To clarify the effects of MGF_360-4L on MDA5 and interferon signaling, we constructed an MGF_360-4L-deficient strain, ASFV/CN/SC/2019-△MGF_360-4L, using homologous recombination and ASFV/CN/SC/2019 as the parental strain (Fig.3D, Fig. S2). The replication ability of ASFV-△MGF_360-4L was slightly lower than that of ASFV-WT (Fig.3E-F). Subsequently, porcine alveolar macrophages (PAMs)were infected with ASFV-WT or ASFV-△MGF_360-4L at different multiplicity of infection (MOIs) (0.1, 1, 2) and collected at 24 hpi. The Western blot results demonstrated that the deletion of MGF_360-4L significantly diminished the suppressive effect of MGF_360-4L on MDA5 protein levels and substantially decreased the inhibition of IRF3 and TBK1 phosphorylation (Fig.3G). These results confirm that ASFV MGF_360-4L may act as an interferon response inhibitory protein to suppress the innate immune response of the host.

**Fig 3.**
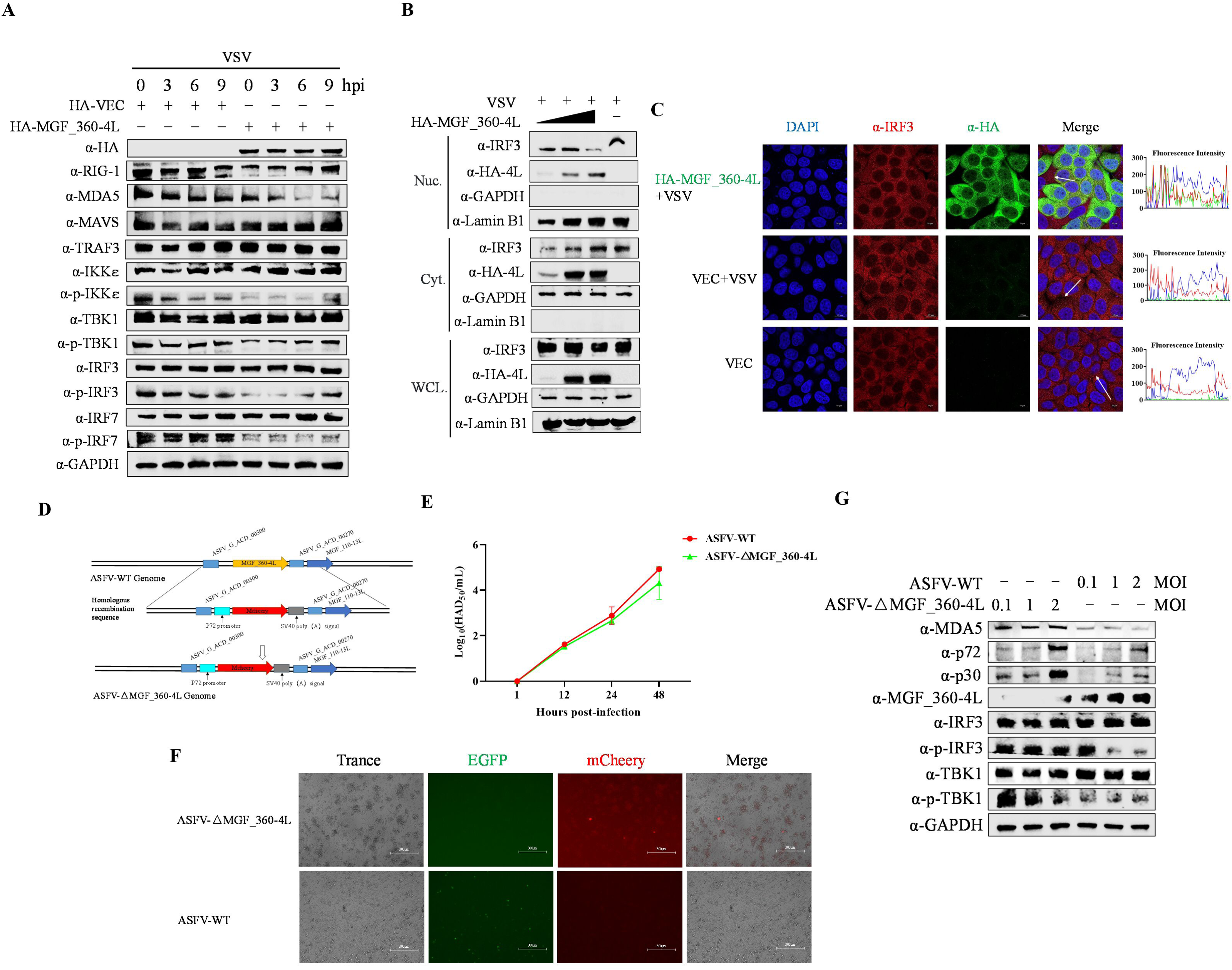
MGF_360-4L inhibits RIG-I/MDA5-mediated interferon signal transduction and suppresses the nuclear translocation of IRF3. (A) HEK293T cells were transfected with pHA-MGF_360-4L following VSV infection for the indicated times. The samples were then collected for protein detection. (B) HEK293T cells were transfected with increasing doses of pHA-MGF_360-4L following VSV infection. The cells were collected and processed for nuclear separation according to the manufacturer’s instructions. The protein expression levels in the nucleus and cytoplasm were subsequently assessed, with GAPDH used as a marker for cytoplasmic components and Lamin B1 used as a marker for the nucleus. (C) HeLa cells were transfected with pHA-MGF_360-4L or empty vector and exposed to VSV or not. The cells were stained with anti-HA and anti-IRF3 antibodies, and their colocalization was observed via confocal microscopy. The fluorescence intensities were determined within the region of interest. (D) Schematic diagram of the construction of the MGF_360-4L deletion mutant. (E-F) PAMs were infected with either ASFV-WT or ASFV-ΔMGF_360-4L (MOI = 3), and the cells were collected at 1, 12, 24, and 48 hours post infection. The viral title was assessed via the HAD_50_ assay (E), and the fluorescence intensities of ASFV-WT carrying the EGFP tag and ASFV-ΔMGF_360-4L carrying the mCherry tag were determined using a fluorescence microscope (F). (G) PAMs were infected with different MOIs (0.1, 1, 2) of ASFV-WT or ASFV-ΔMGF_360-4L for 24 hpi, and cell samples were collected and subjected to Western blotting to assess the phosphorylation levels of IRF3 and TBK1.

### MGF_360-4L induces the degradation of MDA5 via the autophagy‒lysosome pathway

As we described above, MGF_360-4L is known to suppress MDA5 expression, and the potential mechanism(s) underlying MGF_360-4L function were further evaluated. We initially assessed the effect of pHA-MGF_360-4L on pFlag-MDA5. The results indicated that pHA-MGF_360-4L inhibited pFlag-MDA5 protein levels (Fig.4A), but not MDA5 transcript levels in a dose-dependent manner (Fig.4B-C). To further investigate the effects of MGF_360-4L on MDA5 protein levels, the cells were transfected with pFlag-MDA5 with or without pHA-MGF_360-4L or pHA-Vec and then treated with various inhibitors, including the lysosome inhibitor ammonium chloride (NH_4_Cl), the proteasome inhibitor MG-132, the autophagy inhibitor 3-methyladenine (3-MA) or the apoptosis inhibitor Z-VAD-FMK. The Western blot results revealed that NH_4_Cl and 3-MA significantly inhibited the MGF_360-4L-induced degradation of the MDA5 protein, whereas MG-132 and Z-VAD-FMK did not have such effects, suggesting that MGF_360-4L promoted the autophagic degradation of MDA5 (Fig.4D). Additionally, we analyzed the impacts of MGF_360-4L on the autophagy markers LC3 and p62. The Western blot results demonstrated that MGF_360-4L promoted LC3 cleavage and p62 degradation (Fig.4E). Confocal microscopy revealed colocalization of mCherry-LC3B and EGFP-LC3B colocalized primarily around the nucleus in pHA-MGF_360-4L-transfected cells (Fig.4F). In contrast, mCherry-LC3B and EGFP-LC3B were diffusely distributed in the cytoplasm in the absence of pHA-MGF_360-4L (Fig.4F). We further used the autophagy inhibitor chloroquine (CQ) to evaluate the impact of autophagy inhibition on the interferon signaling pathway. The inhibitory effect of MGF_360-4L on MDA5 protein levels was significantly reduced in the presence of CQ, weakening the suppressive effect of MGF_360-4L on interferon signaling (Fig.4G). In summary, these results indicate that MGF_360-4L promoted MDA5 degradation via autophagy, thereby inhibiting interferon signaling.

**Fig 4.**
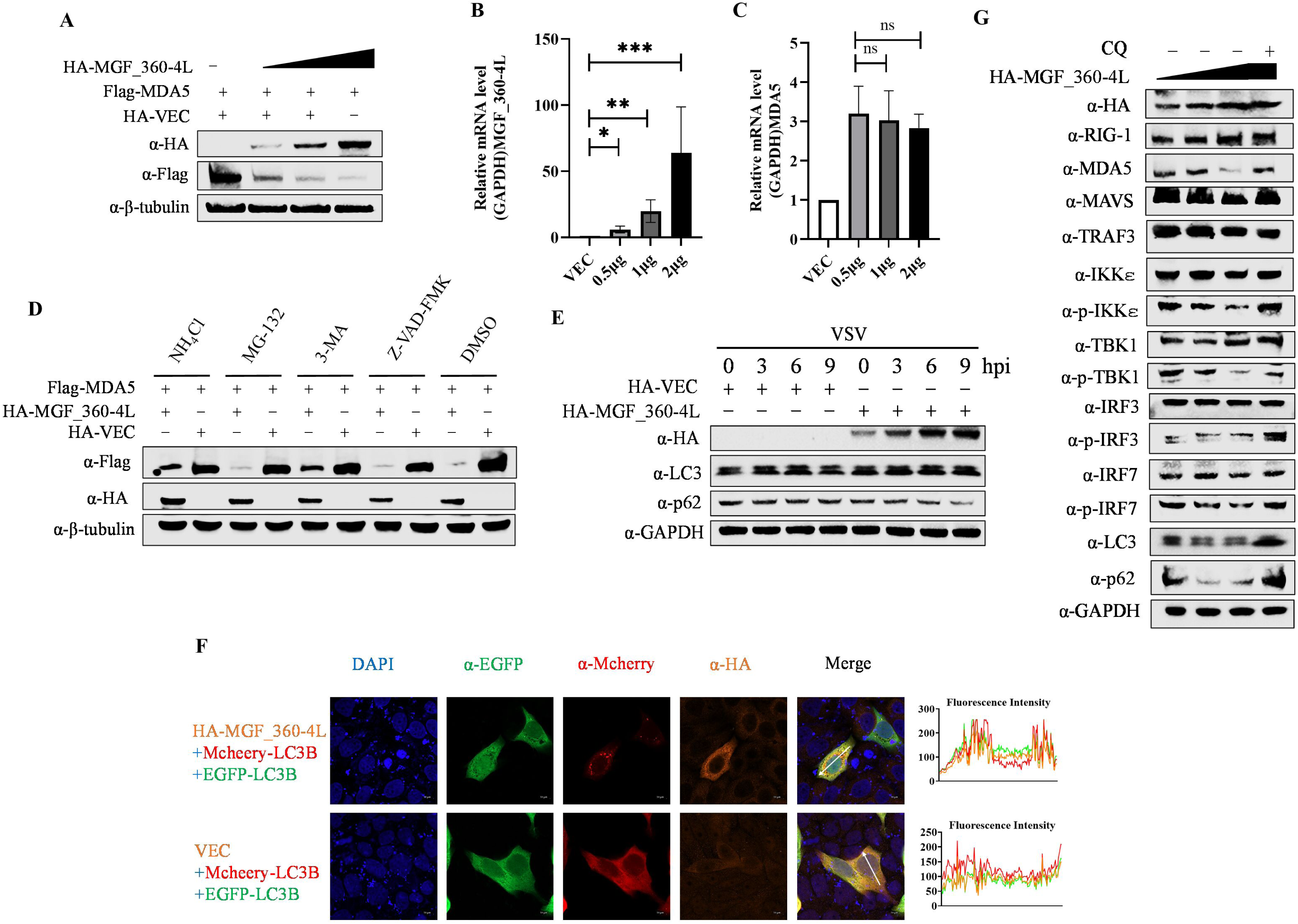
MGF_360-4L promotes the autophagic degradation of MDA5. (A) HEK293T cells were transfected with increasing doses of pHA-MGF_360-4L and pFlag-MDA5. The cells were then collected and subjected to Western blotting with the indicated antibodies to analyze the effect of MGF_360-4L on MDA5 expression. (B-C). HEK293T cells were transfected with increasing doses of pHA-MGF_360-4L, and the transcript levels of MGF_360-4L (B) and MDA5 (C) were measured via RT‒qPCR. (D) HEK293T cells were cotransfected with pFlag-MDA5 and with or without pHA-MGF_360-4L, followed by treatment with DMSO (negative control), NH_4_Cl (20 mM), MG-132 (10 μM), 3-MA (10 mM), or Z-VAD-FMK (20 μM) for 24 h. Then, the cells were collected, and the protein expression levels of MDA5 and MGF_360-4L were analyzed via Western blotting. (E) HEK293T cells were transfected with pHA-MGF_360-4L or pHA-VEC and infected with VSV at the indicated time points. Cell samples were collected, and the protein expression levels of LC3 and P62 were assessed via Western blotting. (F) HeLa cells were cotransfected with Mcheey-LC3B and EGFP-LC3B with or without pHA-MGF_360-4L. The colocalization of pHA-MGF_360-4L and Mcheey-LC3B was observed via fluorescence microscopy. Scale bars, 10 μm. Fluorescence intensity profiles were measured along the drawn measurement line. (G) HEK293T cells were transfected with increasing doses of pHA-MGF_360-4L, followed by treatment with or without the autophagy inhibitor CQ (50 μM). Protein expression was assessed by Western blotting.

### MGF_360-4L recruits the selective autophagy receptor SQSTM1 to induce mitophagy

Our laser confocal microscopy results revealed that MGF_360-4L increased the number of LCB3-positive puncta around the nucleus (Fig.4F). Autophagy occurs through classical and selective autophagy pathways, with selective autophagy requiring the involvement of selective autophagy receptors (28). We further utilized transmission electron microscopy to examine the autophagy induced by MGF_360-4L. The results indicated that MGF_360-4L effectively induced autophagy in cells, with CCCP (carbonyl cyanide 3-chlorophenylhydrazone) used as a positive control for autophagy induction (Fig.5A). Notably, the overexpression of pHA-MGF_360-4L significantly affected mitochondrial morphology, including the disappearance of mitochondrial cristae and alterations in mitochondrial structure, with the presence of autophagic vesicles observed within the mitochondria (Fig.5A). Thus, we hypothesized that MGF_360-4L could induce mitophagy. Mitophagy can occur via two pathways: the NIX/BNIP3 ubiquitin-independent pathway and the PARKIN/PINK1 ubiquitin-dependent pathway (39,40). We thus subsequently assessed the expression levels of the outer mitochondrial membrane protein TOM20 and the mitochondrial matrix proteins HSP60 and VDAC1 following pHA-MGF_360-4L overexpression, with the mitochondrial fission inhibitor Mdivi-1 used as a control. The Western blot results demonstrated that the overexpression of pHA-MGF_360-4L significantly reduced the protein levels of TOM20 and VDAC1, promoted PARKIN expression, and inhibited PINK1 degradation but did not affect the protein levels of NIX and NDP52 (Fig.5B). Additionally, we validated these findings by infecting PAM cells with ASFV-WT or ASFV-△MGF_360-4L at different MOIs (0.1, 1, 2). The Western blot results indicated that the absence of MGF_360-4L stabilized the protein levels of HSP60, VDAC1 and TOM20, maintaining PINK1 degradation (Fig.5C). Therefore, MGF_360-4L induces mitophagy through a selective receptor-PARKIN/PINK1-mediated ubiquitin-dependent pathway.

**Fig 5.**
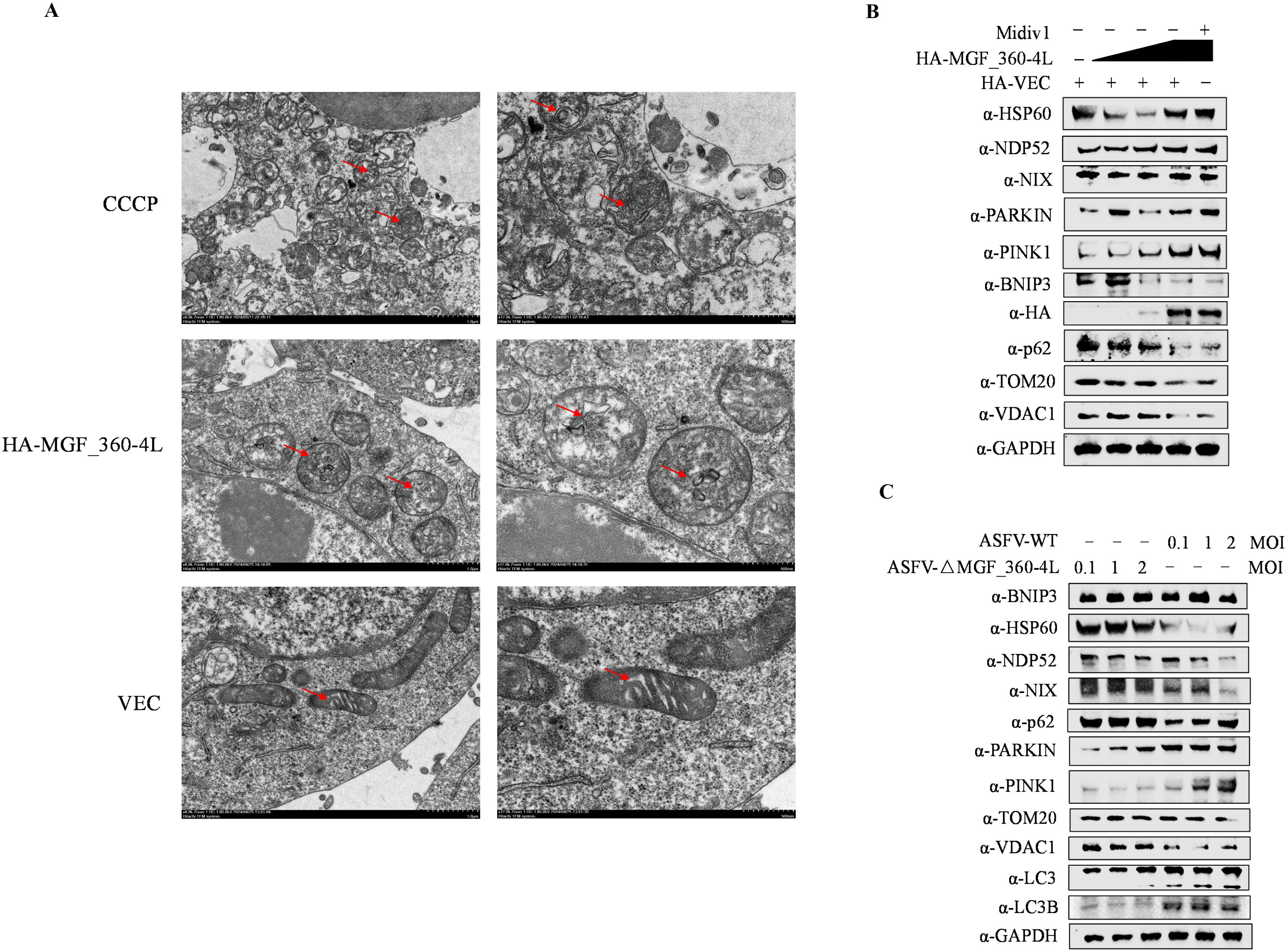
MGF_360-4L induces mitochondrial autophagy. (A) HEK293T cells were transfected with pHA-MGF_360-4L or empty vector. After 20 hours, control cells were treated with the autophagy inducer CCCP for 3 hours. Ultrathin sections of cellular structures were observed via electron microscopy, and autophagic vacuoles are indicated (red arrows). (B) HEK293T cells were transfected with increasing doses of pHA-MGF_360-4L or pHA-VEC, followed by treatment with or without Midiv 1 (mitochondrial division inhibitor, 1,20 μM). The cells were collected, and the protein expression levels were detected via Western blotting. (C). PAMs were infected with different MOIs (0.1, 1, 2) of ASFV-WT or ASFV-ΔMGF_360-4L for 24 hpi. The cells were collected, and the protein expression levels were assessed via Western blotting.

To further clarify that MGF_360-4L uses a specific autophagy receptor to induce autophagy, the cells were cotransfected with pHA-MGF_360-4L with or without pMyc-OPTN, pMyc-NBR1, pMyc-TOLLIP, pMyc-NDP52, pMyc-SQSTM1/p62, or pMyc-VEC, and subsequent Co-IP experiments revealed that MGF_360-4L specifically interacts with p62 (Fig.6A-B). GST pull-down assays further confirmed that MGF_360-4L directly interacts with p62 (Fig.6C). Laser confocal microscopy revealed that MGF_360-4L colocalized with p62 around the nucleus (Fig.6D-E) but did not colocalize with NDP52 (Fig.6F). Thus, we hypothesize that p62 is the primary selective autophagy receptor through which MGF_360-4L induces the autophagic degradation of MDA5.

**Fig 6.**
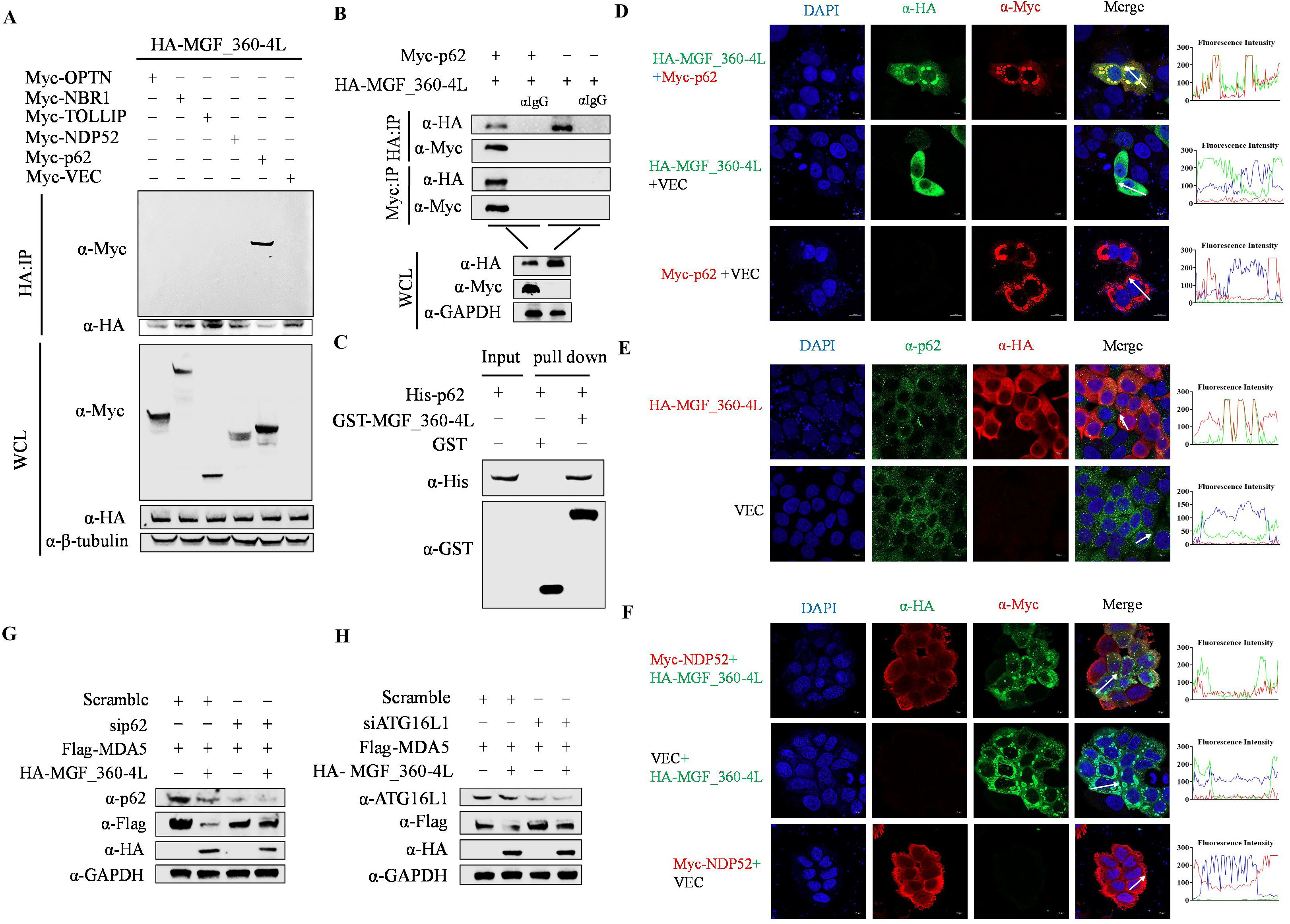
MGF_360-4L recruits the selective autophagy receptor p62 to induce mitophagy. (A-B) HEK293T cells were cotransfected with pHA-MGF_360-4L and Myc-OPTN, Myc-NBR1, Myc-TOLLIP, Myc-NDP52, or Myc-p62, and the interactions between proteins were detected by Co-IP. (C) GST-tagged MGF_360-4L protein (10 μg) or GST antigen was added to GST-Sepharose 4B beads, which were incubated for 3 h at room temperature, followed by binding with His-tagged p62 protein (10 μg). The samples were subjected to Western blotting analysis to detect direct interactions between MGF_360-4L-GST and p62-His. (D-E) HEK293T cells were transfected with pFlag-MDA5 with or without pHA-MGF_360-4L, followed by the addition of siRNA targeting p62 or ATG16L1 or a scramble siRNA as a control. The cells were collected, and the protein expression levels were analyzed via Western blotting. (F-H) HeLa cells were transfected with pHA-MGF_360-4L and pMyc-p62, NDP52 or empty vector. The colocalization of MGF_360-4L with p62 or NDP52 was detected via confocal microscopy. Fluorescence intensity profiles were measured along the drawn measurement line.

Conversely, an analysis of the lysates of cells cotransfected with p62-directed siRNA and a plasmid encoding Flag MDA5 with or without HA-MGF_360-4L revealed that p62 silencing inhibited MGF_360-4L-induced degradation of MDA5 (Fig.6G). Additionally, we screened autophagy-related genes (ATGs) for their role in MGF_360-4L-mediated MDA5 degradation. The Western blot results indicated that depletion of ATG16L1 blocked the MGF_360-4L-induced degradation of MDA5 but not that of ATG5, ATG7 or ATG12 (Fig.6H, Fig. S3A-C). These results demonstrate that MGF_360-4L recruits p62 to induce mitophagy and degrade MDA5 through the ATG16L1-dependent PARKIN/PINK1 pathway.

### MGF_360-4L affects the interaction between MDA5 and MAVS and impairs ISG15-mediated ISGylation and activation of MDA5

During viral infection, dsRNA produced by the virus is detected by the MDA5 receptor, which then recruits MAVS/VISA to transmit signals and activate the interferon signaling pathway (20). In addition to being activated through the detection of dsRNA, MDA5 is also activated via ISG15-mediated ISGylation, which contributes to its antiviral functions. (26). We previously reported that MGF_360-4L greatly increases ISG15 transcript levels, with a more pronounced effect in the presence of IFN-β (Fig.1F). Thus, to elucidate the role of ISG15 in the effect of MGF_360-4L on MDA5, we cotransfected pHA-MAVS, pFlag-MDA5 and increasing doses of pMyc-MGF_360-4L, along with treatment with CQ to ensure MDA5 stability. Co-IP results demonstrated that MGF_360-4L inhibits the interaction between MDA5 and MAVS in a dose-dependent manner (Fig.7A). Furthermore, the results of laser confocal microscopy revealed that in the presence of MGF_360-4L, MDA5 colocalizes with MAVS and anchors at the mitochondria, whereas in the absence of MGF_360-4L, MDA5 is distributed throughout the cytoplasm (Fig.7B). To assess the ISGylation of MDA5, the cells were cotransfected with pFlag-MDA5, pMyc-ISG15, or increasing doses of pHA-MGF_360-4L and treated with CQ to ensure MDA5 stability. Co-IP results indicated that MGF_360-4L inhibits the ISGylation of MDA5 induced by ISG15 in a dose-dependent manner, whereas MGF_360-4L itself does not undergo ISGylation (Fig.7C). The results of laser confocal microscopy further revealed that MDA5 and ISG15 can colocalize in the cytoplasm (Fig.7D). Collectively, these findings demonstrate that MGF_360-4L inhibits the interaction between MDA5 and MAVS/VISA and impairs ISG15-mediated ISGylation and activation of MDA5.

**Fig 7.**
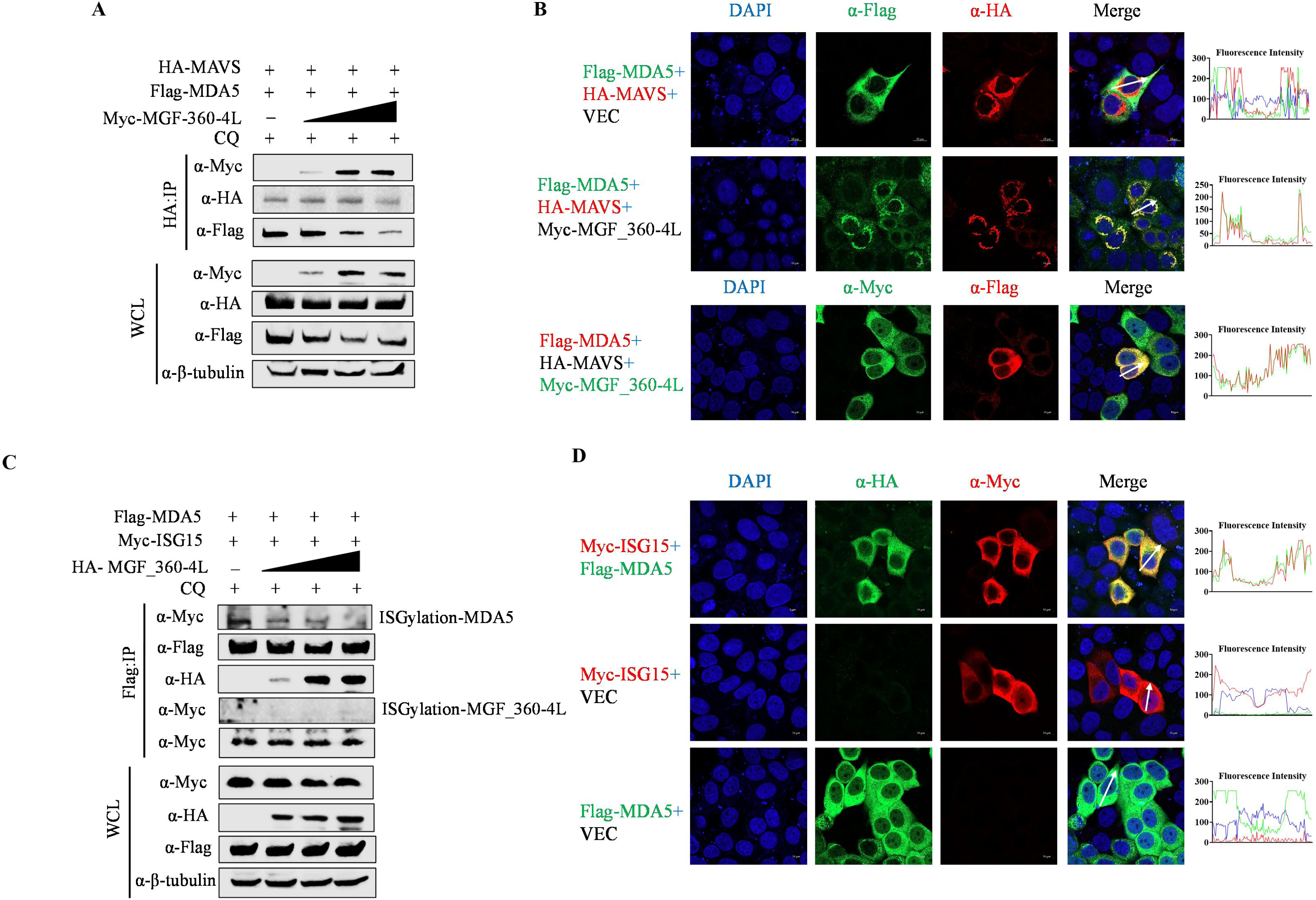
MGF_360-4L inhibits the interaction between MDA5 and MAVS/VISA and reduces the ISGylation activation of MDA5 by ISG15. (A) HEK293T cells were transfected with increasing doses of pMyc-MGF_360-4L along with pHA-MAVS and pFlag-MDA5. CQ (50 μM) was added to ensure the stability of MDA5. After 24 hours, the cell samples were collected, and the interactions between proteins were detected via Co-IP. (B) HeLa cells were transfected with the indicated plasmids. The colocalization of pMyc-MGF_360-4L with pHA-MAVS and pFlag-MDA5 was detected via confocal microscopy. Fluorescence intensity profiles were measured along the drawn measurement line. (C) HEK293T cells were transfected with increasing doses of the pHA-MGF_360-4L plasmid along with the pFlag-MDA5 and pMyc-ISG15 plasmids. CQ (50 μM) was added to ensure the stability of MDA5. After 24 hours, the cell samples were collected, and the protein interactions and ISGylation levels of MDA5 and MGF_360-4L were assessed via Co-IP. (D) HeLa cells were transfected with pFlag-MDA5 and pMyc-ISG15 or with an empty vector as a control. The colocalization of ISG15 and MDA5 was detected via confocal microscopy. Fluorescence intensity profiles were measured along the drawn measurement line.

### MGF_360-4L is ubiquitinated and degraded by OAS1

Our previous Co-IP/LC‒MS/MS data revealed that OAS1 is predicted to interact with MGF_360-4L. Therefore, we constructed a pFlag-OAS1 plasmid to evaluate its function in MGF_360-4L. First, we confirmed the interaction between MGF_360-4L and OAS1 via Co-IP (Fig.8A). GST-pull down assays also demonstrated that MGF_360-4L directly interacts with OAS1 (Fig.8B). We subsequently assessed the effect of OAS1 on MGF_360-4L, and cells were cotransfected with pMyc-MGF_360-4L and increasing doses of pFlag-OAS1. The Western blot results revealed that OAS1 inhibited the protein expression of MGF_360-4L in a dose-dependent manner (Fig.8C). Similarly, a degradation pathway screening assay using inhibitors such as MG-132, 3-MA, and NH_4_Cl indicated that OAS1 degrades MGF_360-4L via the ubiquitin‒proteasome pathway (Fig.8D). We then constructed plasmids with six different ubiquitin mutation sites (K6, K11, K27, K29, K33, K48, and K63) for ubiquitination site validation. Co-IP revealed that OAS1 ubiquitinates MGF_360-4L through polyubiquitin chains linked via K6, K33, K48, and K63 (Fig.8E). To further identify the main ubiquitination sites on MGF_360-4L, we performed ubiquitination assays on the aforementioned MGF_360-4L fragments. Co-IP results revealed that the ubiquitination level of the MGF_360-4L-3 fragment was significantly lower than that of the other MGF_360-4L fragments (Fig.8F). Thus, we hypothesize that the primary ubiquitination sites on MGF_360-4L are in the Alpha-lactotoxin LT1a-like domain. Furthermore, we performed site-specific mutagenesis on lysine (K) residues in the Alpha-lactotoxin LT1a-like domain, constructing ten Flag-tagged MGF_360-4L mutant plasmids (201R, 208R, 209R, 257R, 290R, 295R, 320R, 321R, 327R, and 329R). Co-IP results indicated that the ubiquitination levels of MGF_360-4L were significantly reduced following the mutation of lysine residues 209, 295, and 327 (Fig.8G). In summary, OAS1 ubiquitinates and degrades MGF_360-4L through polyubiquitin chains linked via K6, K33, K48, and K63, and lysine residues 209, 295 and 327 of MGF_360-4L are crucial sites for the ubiquitination of MGF_360-4L.

**Fig 8.**
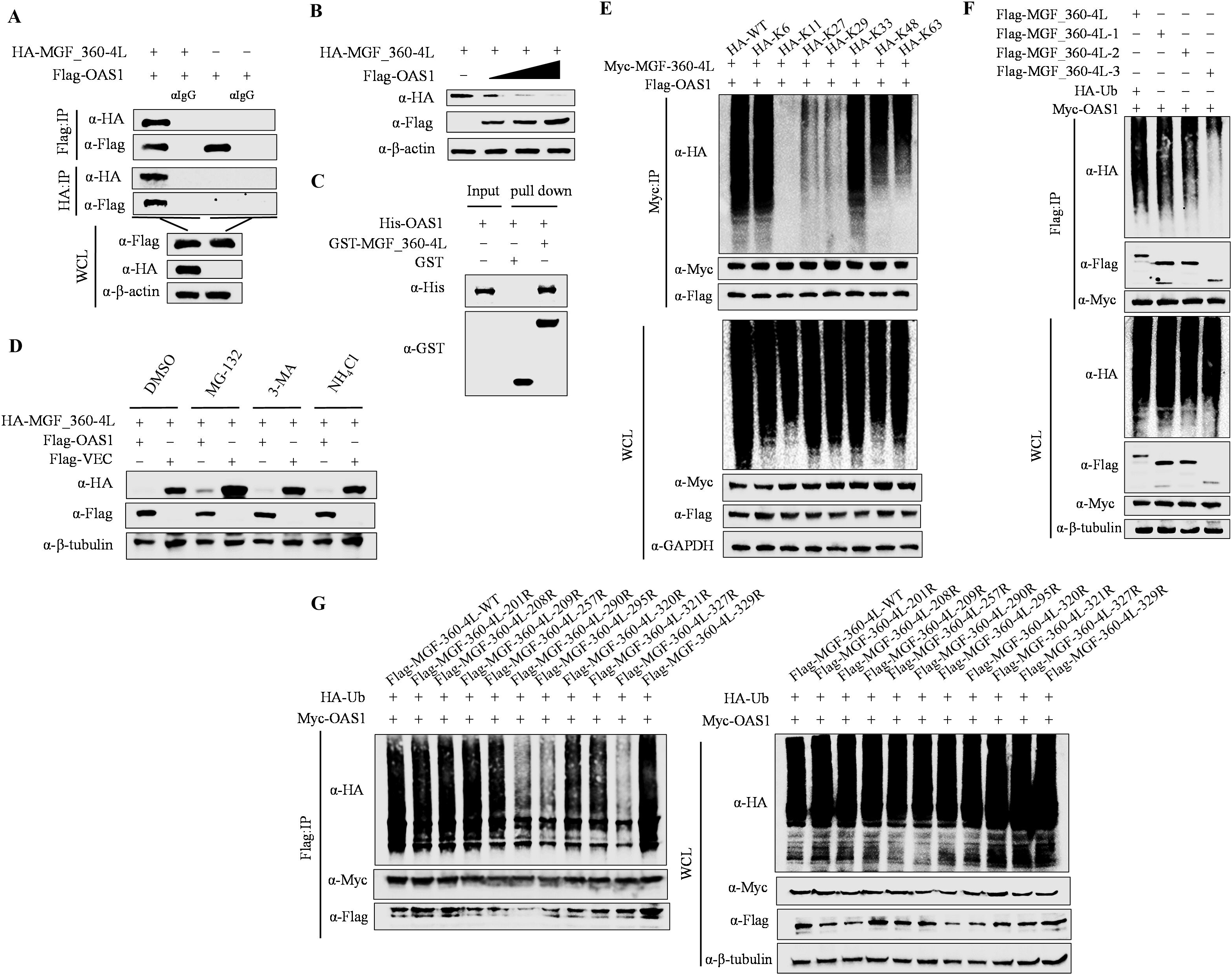
OAS1 degrades MGF_360-4L through polyubiquitination, with K290, K295, and K327 as primary ubiquitination sites. (A) HEK293T cells were transfected with pFlag-OAS1 and pHA-MGF_360-4L. After 24 hours, the cell samples were collected, and the interaction between OAS1 and MGF_360-4L was assessed via Co-IP. (B) HEK293T cells were transfected with increasing doses of pFlag-OAS1 and pHA-MGF_360-4L. After 24 hours, the cell samples were collected, and the protein expression levels were analyzed via Western blotting. (C) GST-tagged MGF_360-4L protein (10 μg) or GST antigen was added to GST-Sepharose 4B beads, which were incubated for 3 h at room temperature, followed by binding with His-tagged OAS1 protein (10 μg). The samples were subjected to Western blotting analysis to detect direct interactions between MGF_360-4L-GST and OAS1-His. (D) HEK293T cells were cotransfected with pHA-MGF_360-4L with or without pFlag-OAS1. After 20 hours, the cells were treated with NH_4_Cl (20 mM), MG132 (10 μM), 3-MA (10 mM), or DMSO as a control. Protein expression was analyzed by Western blotting. (E) HEK293T cells were transfected with pMyc-MGF_360-4L and pFlag-OAS1 plasmids, along with six ubiquitin retention mutant plasmids (K6, K11, K27, K29, K33, K48, and K63). After 24 hours, the cells were collected, and the degree of MGF_360-4L ubiquitination was assessed via Co-IP. (F) HEK293T cells were transfected with pFlag-MGF_360-4L 1-3 domains along with pMyc-OAS1 or HA-Ub. After 24 hours, the cells were collected, and the ubiquitination levels of MGF_360-4L were analyzed via Co-IP. (G) HEK293T cells were transfected with pMyc-OAS1 and HA-Ub and ten point mutation plasmids (201R, 208R, 209R, 257R, 290R, 295R, 320R, 321R, 327R, and 329R) based on lysine residues in the Alpha-lactotoxin LT1a-like domain (MGF_360-4L-3) of MGF_360-4L. After 24 hours, the cells were collected, and the primary ubiquitination sites of MGF_360-4L were identified via Co-IP.

### ASFV-△MGF_360-4L significantly reduces the virulence of the parental strain and increases resistance to lethal challenge in pigs

To evaluate the pathogenicity of ASFV-△MGF_360-4L in pigs, we conducted a challenge experiment. Pigs were injected intramuscularly with ASFV-△MGF_360-4L (20 HAD_50_, n=5), ASFV-△MGF_360-4L (100 HAD_50_, n=5), or ASFV-WT (20 HAD_50_, n=5), with a control group receiving the same volume of RPMI 1640 (n=5). Disease progression was monitored for 21 days. Pigs injected with ASFV-WT (20 HAD_50_) presented a fever at 3 days postinfection (dpi), with temperatures reaching 41.7°C, which persisted until 7 dpi and then decreased (Fig.9A); however, 3 pigs died at 8 dpi, and 2 pigs died at 9 dpi (Fig.9A and B). In contrast, pigs in the ASFV-△MGF_360-4L (20 HAD_50_) group presented a slight temperature increase (≤41°C) postinoculation, which returned to normal. No significant clinical signs of ASF were observed, and all pigs in this group survived. The control group presented normal temperatures and no deaths over the 21-day period. Pigs in the ASFV-△MGF_360-4L (100 HAD_50_) group also presented a slight increase in temperature (≤41°C), which subsequently normalized, with all pigs surviving. At 21 dpi, these pigs were challenged with ASFV-WT (20 HAD_50_), and some pigs presented transient fever but recovered without showing clinical signs of ASF and did not die by 23 dpi (Fig.9A and B). At 3 dpi, ASFV-WT (20 HAD_50_) was detected in swabs and blood samples, with more viral genome copies than the other groups, reaching peak levels just before death (8–9 dpi) (Fig.9C and 9D). Moreover, viral genome copies in the blood and swabs from the ASFV-△MGF_360-4L (100 HAD_50_) and ASFV-△MGF_360-4L (20 HAD_50_) groups were almost completely cleared by 16–48 dpi (Fig.9C and D). The viral load in the organs/tissues from the ASFV-WT group was significantly greater than that in the organs/tissues from the ASFV-△MGF_360-4L group (Fig.9E). The heart, liver, spleen, lungs, kidneys, tonsils, mandibular lymph nodes, inguinal lymph nodes, and mesenteric lymph nodes from pigs challenged with ASFV-WT (20HAD_50_) presented hemorrhage, congestion, and edema. In contrast, pigs challenged with ASFV-△MGF_360-4L (100 HAD_50_) presented no obvious lesions; similarly, ASF-specific pathological changes were almost undetectable in the ASFV-△MGF_360-4L (20 HAD_50_) group (Fig.9F).

**Fig 9.**
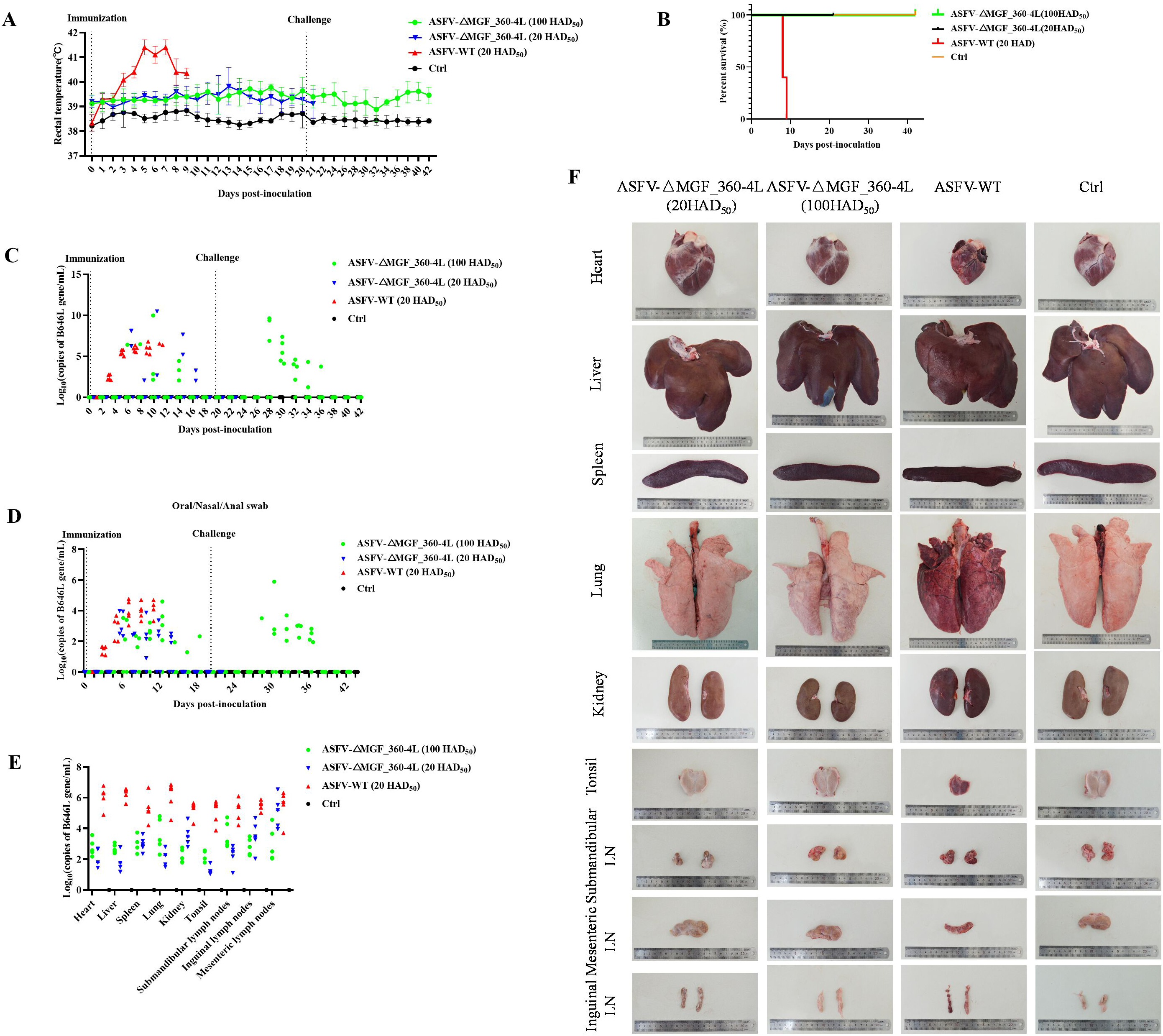
MGF_360-4L is a critical virulence gene of ASFV. (A) Rectal temperatures of pigs were measured daily after intramuscular injection of ASFV-WT (20 HAD_50_) or ASFV-ΔMGF_360-4L (20 HAD_50_, 100 HAD_50_) (n=5). (B) Survival rates of pigs were calculated. (C-D) qPCR analysis of ASFV genomic DNA copy number in blood, oral, nasal, and anal swabs obtained from pigs after infection with either ASFV-ΔMGF_360-4L or ASFV-WT. (E) qPCR analysis of ASFV genomic DNA copy number in various organs (heart, liver, spleen, lung, kidney, submandibular lymph nodes, mesenteric lymph nodes, inguinal lymph nodes, and tonsils) from pigs infected with ASFV-ΔMGF_360-4L or ASFV-WT.

Histopathological examination revealed hemorrhage, congestion, and edema in the heart, liver, spleen, lungs, kidneys, tonsils, mandibular lymph nodes, inguinal lymph nodes, and mesenteric lymph nodes of ASFV-WT (20 HAD_50_)-infected pigs. In contrast, pigs challenged with ASFV-△MGF_360-4L (100 HAD_50_) presented only mild lesions, with minor congestion and hemorrhage in the lungs and mandibular lymph nodes. Additionally, no significant pathological changes were observed in pigs infected with ASFV-△MGF_360-4L (20 HAD_50_). In addition to a few hemorrhagic spots in the interstitial muscle tissue of the heart, extensive diffuse hemorrhage and inflammatory cell infiltration were observed in the liver, spleen, lungs, kidneys, tonsils, and lymph nodes (including the submandibular, inguinal, and mesenteric lymph nodes) in the ASFV-WT group. Additionally, substantial tissue fluid exudation was observed in the kidneys and lungs. In contrast, the pigs inoculated with ASFV-△MGF_360-4L (20 HAD_50_) or ASFV-△MGF_360-4L (100 HAD_50_) presented only mild histopathological changes, with minimal hemorrhage and tissue fluid exudation observed (Fig.10A). The ELISA results revealed that antibodies were detected in the serum of pigs challenged with ASFV-△MGF_360-4L (20 HAD_50_ and 100 HAD_50_), and antibodies were also detectable in the ASFV-△MGF_360-4L (100 HAD_50_) group following challenge with the parental virus (Fig.10B). Furthermore, higher levels of IFN-β were detected in the serum of pigs in the ASFV-△MGF_360-4L group (20 HAD_50_ and 100 HAD_50_) than in the ASFV-WT group (Fig.10C). Overall, our results demonstrate that MGF_360-4L is a virulence gene and that ASFV-△MGF_360-4L reduces pathogenicity and significantly weakens viral replication capacity in pigs, enabling resistance to challenge by the parental ASFV-WT strain.

**Fig 10.**
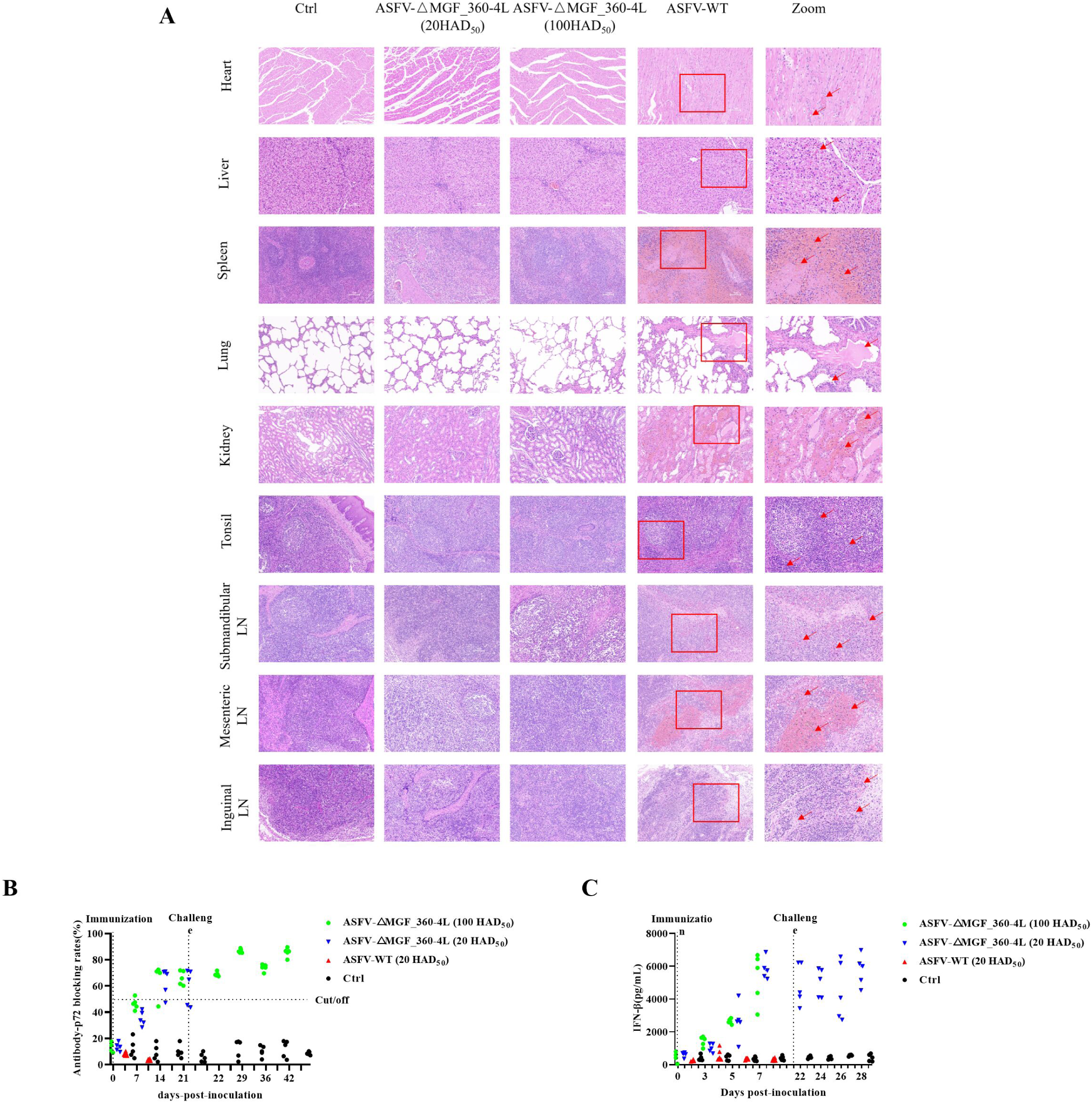
Compared with ASFV-WT, ASFV-ΔMGF_360-4L attenuated the pathogenicity of ASFV-WT and induced greater antibody responses and interferon levels. (**A**) Hematoxylin and eosin staining was used to assess the pathological changes caused by ASFV-△MGF_360-4L and ASFV-WT in different pig tissues (red arrows in the figure). Scale bar = 50 μm. **(B)** The levels of antibodies against p72 were measured in pigs vaccinated with ASFV-ΔMGF_360-4L (20 HAD_50_, 100 HAD_50_) or ASFV-WT (20 HAD_50_) using a commercial ELISA kit, and the results are presented as blocking percentages. Antibody positivity was defined as a blocking rate ≥50%, negativity was defined as a blocking rate of ≤40%, and results ranging between 40% and 50% were considered undetermined. **(C)** The levels of IFN-β were measured in pigs vaccinated with ASFV-ΔMGF_360-4L (20 HAD_50_, 100 HAD_50_) or ASFV-WT (20 HAD_50_) using a commercial ELISA kit.

## Discussion

Previous studies have shown that several virulence proteins encoded by ASFV genes, such as DP96R (41), E120R (42), MGF_505-7R (32), MGF_505-11R (43) and pI215L (44), are capable of suppressing host immunity and blocking the activation of the interferon pathway to limit the secretion of IFN-β. Recent reports have indicated that ASFV can hijack host protein degradation mechanisms and induce autophagy to counteract innate immunity (30-34). However, ASFV genes, particularly those in the MGF that affect interferon pathway activation, remain largely unknown. In this study, we identified a novel innate immune suppression gene, MGF_360-4L, which specifically targets the dsRNA receptor MDA5. MGF_360-4L recruits the selective autophagy receptor SQSTM1/p62 to promote mitochondrial autophagy and degrade MDA5, thereby limiting interferon signal transduction and suppressing IFN secretion. Previous research has demonstrated that deletion of MGF 360 genes (9L, 10L, 11L, 12L, 13L, 14L) can reduce ASFV virulence and provide resistance to lethal attacks from the parental strain, but the effects of individual genes on virulence reduction are poorly understood. Gene deletions of MGF_300-2R (30), MGF_505-7R (32), and MGF_360-9L (35) can attenuate virulence in pigs, but whether they can provide protection against lethal attacks by the parental strain is unclear. In our study, ASFV-△MGF_360-4L was generated from the ASFV-WT parental strain and was found to attenuate virulence in pigs and provide protection against lethal attacks from the parental strain. The IFN-β levels in pigs challenged with ASFV-△MGF_360-4L were significantly greater than those in pigs challenged with ASFV-WT (Fig.10B). Observations at the organ/tissue level also revealed minimal tissue damage and pathological changes in pigs inoculated with ASFV-△MGF_360-4L. Our research indicates that MGF_360-4L-induced mitochondrial autophagy represents a novel mechanism by which ASFV evades host immune responses and promotes viral replication.

RIG-I/MDA5, as host cytoplasmic PRRs, play crucial roles in regulating host type I and III antiviral immune responses and the expression of proinflammatory cytokines (45). Additionally, PRRs can be regulated by host-induced autophagy to maintain homeostasis (46). The interferon signaling modulation of RIG-I/MDA5 relies on the formation of the MAVS/VISA complex, which further promotes the phosphorylation of IKK, TBK1, and IRF3 through TRAF3 signaling (20), leading to the nuclear translocation of IRF3 and subsequent increases in IFN secretion and ISG production (47). MDA5 activation can also depend on ISG15-mediated ISGylation of the CARD domain to stimulate innate immune responses (26). Our results revealed that MGF_360-4L decreases the protein level of MDA5 and obstructs interferon signaling (Fig.3A-B). Furthermore, the Alpha-lactotoxin_LT1a-like domain of MGF_360-4L interacts specifically with the SF2_C dicer domain of MDA5 without interacting with RIG-I (Fig.2F-G). This specific interaction limits the association between MDA5 and MAVS/VISA (Fig.7A). Moreover, IFN-induced ISGs can inhibit viral infection, with ISG15 being particularly notable. ISG15-induced ISGylation occurs in the modification processes of both host and viral proteins (48). Suppression of STAT1 ISGylation can exacerbate hepatitis C virus (HCV) infection (49), whereas ISGylation of IRF3 can weaken its interaction with PIN1, maintaining IRF3 activation and thereby inhibiting SeV replication (50). Previous studies have also indicated that ISGylation of the IAV NS1 protein can restrict its nuclear translocation and enhance the host antiviral response (51). ISGylation of HPV-16 capsid proteins can prevent the release of mature viral particles, reducing infectivity (52). Our results demonstrate that overexpression of MGF_360-4L significantly stimulates the transcription of ISG15 and works synergistically with IFN-β while inhibiting the transcription of IFIT1, IFIT2, OASL, STAT1 and STAT2 (Fig.1F). However, MGF_360-4L hinders MDA5 ISGylation without causing ISGylation of MGF_360-4L itself (Fig.7C). This specific induction of ISG15 transcription may further influence the ISGylation of host or viral proteins, thereby affecting the host’s innate immune response against viral infections. However, these specific mechanisms need further investigation.

Autophagy is a crucial pathway in eukaryotes for the clearance of misfolded proteins and damaged organelles, contributing to the maintenance of cellular metabolic homeostasis in the host (53). During autophagy, selective autophagy receptors such as SQSTM1/p62, OPTN and NBR1 recruit LC3 through the LC3 interaction region (LIR), promoting the formation and expansion of autophagosomes, which ultimately fuse with lysosomes to form autolysosomes and are subsequently degraded (54,55). Mitophagy, a specific form of autophagy, is essential for removing damaged mitochondria and occurs via classical and nonclassical pathways. The classical pathway relies on the ubiquitin-dependent PARKIN-PINK1 pathway (40,56). When mitochondria are damaged, they are marked by PINK1, which then recruits the E3 ubiquitin ligase PARKIN to accumulate on the mitochondria, inducing mitophagy (57). Additionally, NIX and BNIP3 can also induce mitophagy through the nonclassical pathway (58). Previous studies have demonstrated that ASFV infection can induce autophagy (30-34), but there are few reports on whether ASFV proteins can induce mitophagy. Our results showed that MGF_360-4L promoted the formation of EGFP-LC3B and mCherry-LC3B puncta and their colocalization around the nucleus, which demonstrated that MGF_360-4L induced autophagy (Fig.4F). The autophagy inhibitor CQ reduced the effect of MGF_360-4L on MDA5 degradation and restored interferon signaling (Fig.4G). Furthermore, MGF_360-4L induced mitophagy through the PARKIN-PINK1 ubiquitin-dependent pathway (Fig.5A-C). The maturation of autophagosomes also depends on the ATG12-ATG5-ATG16L1 complex, which is located in the cytosol and preautophagosomal structures (59). Subsequently, autophagosomes interact with LC3 to induce the formation of mature autophagosomes (60). Our results indicate that MGF_360-4L mediates autophagy through the p62 selective receptor rather than NDP52 (Fig.6A-C, F) and primarily relies on the ATG16L1 autophagy-related molecule to regulate the degradation of MDA5 rather than ATG5, ATG7, or ATG12 (Fig.6H, Fig. S2B-D). These findings suggest that MGF_360-4L can degrade MDA5 to inhibit interferon signaling, thereby achieving immune evasion in the host.

In our previous study, we demonstrated that the interferon-stimulated gene OAS1 can inhibit virus maturation and particle assembly and restrict viral replication by recruiting Trim21 to degrade the major structural protein p72 of ASFV (37), making OAS1 a potential host restriction factor against ASFV. However, our Co-IP/LC‒MS/MS data revealed that OAS1 interacts with multiple ASFV viral proteins, including p72, p49, I329L, MGF_360-4L, MGF_360-8L and NP868R (Fig.S1). To further elucidate the role of OAS1 in antiviral defense against ASFV, we explored the interaction between OAS1 and MGF_360-4L and the underlying mechanisms involved. Our results indicate that OAS1 can ubiquitinate and degrade MGF_360-4L, with the lysine residues at positions 209, 257 and 327 being the primary ubiquitination sites (Fig.8A-C, G). These findings provide further insights into the mechanism by which OAS1 inhibits ASFV replication, offering new perspectives on host defense against ASFV. However, whether OAS1 also participates in other aspects of the immune response during ASFV infection remains to be further validated.

In summary, our study revealed that infection with the MGF_360-4L deletion mutant induces increased levels of IFN both *in vivo* and *in vitro*. ASFV-ΔMGF_360-4L immunization provides protection against the lethal attack of ASFV-WT, making it a promising candidate vaccine for ASFV immunity. Mechanistically, MGF_360-4L induces mitophagy and degrades MDA5 through PARKIN-PINK1 ubiquitin-dependent pathway after the recruitment of the selective autophagy receptor p62 while also inhibiting the ISGylation of MDA5 and its interaction with MAVS, thereby limiting interferon signaling. Additionally, OAS1 ubiquitinates and degrades MGF_360-4L at lysine residues 209, 257, and 327, further elucidating the intrinsic mechanism by which OAS1 suppresses ASFV replication (Fig.11). Our findings contribute to the understanding of MGF_360-4L function and its role in viral pathogenesis, offering new insights into ASFV prevention and control.

**Fig 11.**
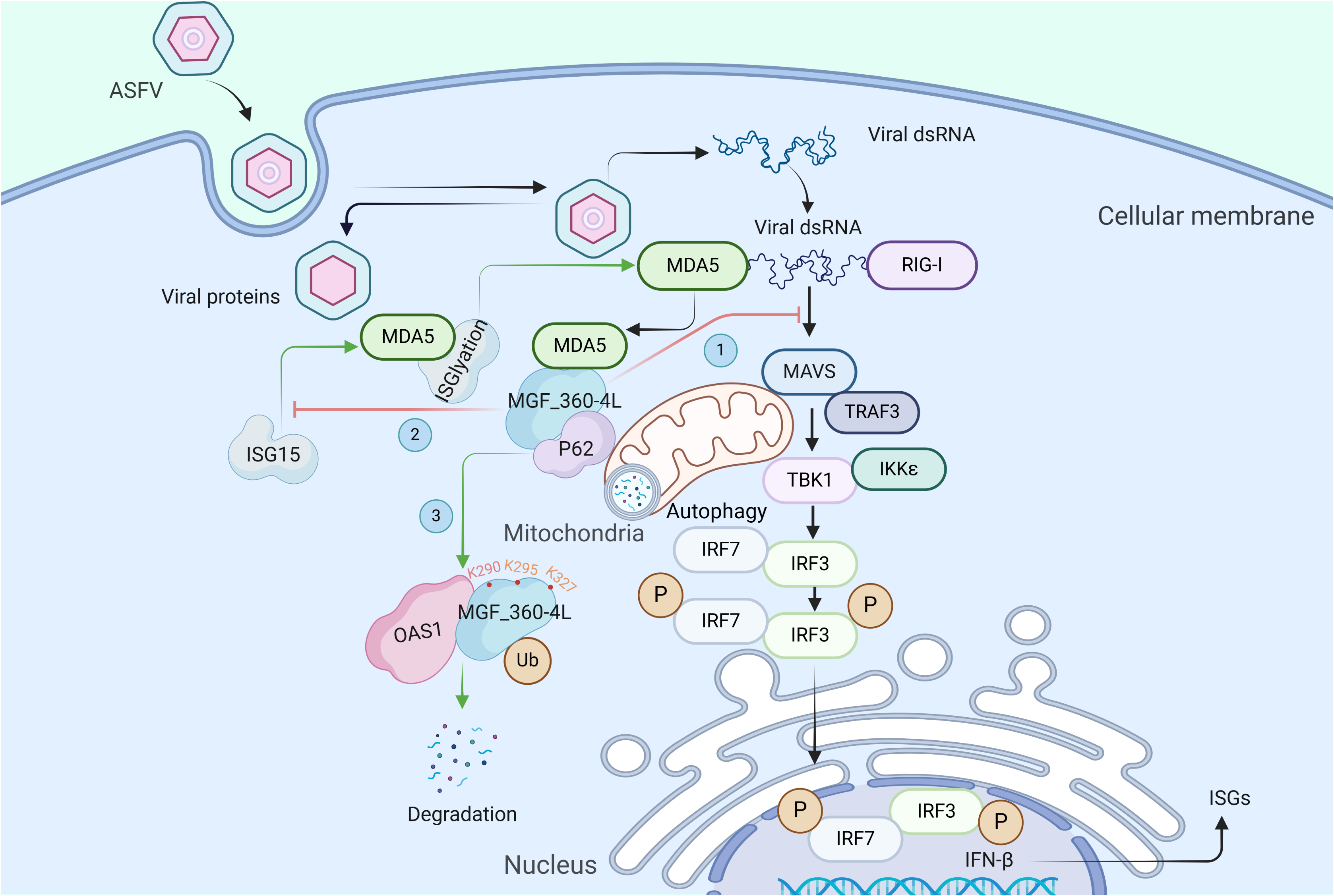
Schematic illustrating the inhibition of interferon signaling by MGF_360-4L-mediated degradation of MDA5 through autophagy. After ASFV enters cells, its dsRNA is sensed by RIG-I/MDA5, which recruits MAVS/VISA to activate interferon signaling. MGF_360-4L can degrade MDA5 through mitochondrial autophagy, thereby impeding the interaction between MDA5 and MAVS/VISA, inhibiting the nuclear translocation of IRF3, and suppressing interferon signaling. ISG15 can activate interferon signaling by ISGylating MDA5 during viral infection. MGF_360-4L can inhibit the ISGylation of MDA5, thereby inhibiting interferon signaling. OAS1 can interact with MGF_360-4L and induce its degradation through polyubiquitination. The primary ubiquitination sites on MGF_360-4L are K290, K295, and K327.

## Materials and Methods

### Biosafety and ethical statement

All experiments involving ASFV were conducted at the Biosafety Level 3 (BSL-3) laboratory of the Lanzhou Veterinary Research Institute (LVRI), Chinese Academy of Agricultural Sciences (CAAS). This laboratory is accredited by the China National Accreditation Service for Conformity Assessment (CNAS) and approved by the Ministry of Agriculture and Rural Affairs. The handling of the animals in this study adhered to the animal ethics procedures and guidelines of LVRI and the Chinese Academy of Agricultural Sciences (Approval number: LVRIAEC-2023-043). To mitigate any potential risks in the laboratory, strict adherence to protocols was enforced, with all activities monitored by professional staff at LVRI. Additionally, local and central government authorities conducted random inspections without prior notification.

### Cells and viruses

The human embryonic kidney cell line HEK293T and the human cervical cancer cell line HeLa were cultured in Dulbecco’s modified Eagle’s medium (DMEM; Gibco, China) supplemented with 10% fetal bovine serum (FBS; Gibco, 16000–044) and 1% penicillin‒ streptomycin (Gibco, 1514012). The immortalized porcine alveolar macrophage line 3D4/21 (CRL-2843) was maintained in RPMI 1640 medium (Gibco, 11875085) supplemented with 10% FBS and 1% penicillin‒streptomycin. Primary PAMs from 30-day-old healthy pigs were cultured in RPMI 1640 medium (Gibco, 11875085) supplemented with 10% FBS and 1% penicillin‒streptomycin. All the cells were grown in a humidified incubator with 5% CO_2_ at 37°C. ASFV CN/SC/2019 was isolated, identified, and maintained in the LVRI-BSL-3 laboratory of the China African Swine Fever Regional Laboratory (Lanzhou). Using ASFV CN/SC/2019 as the parental strain, the recombinant virus ASFV-EGFP with an EGFP-tag insertion was constructed and maintained in the LVRI-BSL-3 facility.

### Antibodies and reagents

Monoclonal antibodies against p72, p30 and MGF_360-4L were generated and stored in our laboratory. The following antibodies were purchased from Cell Signaling Technology (USA): rabbit anti-DYKDDDK-Tag (CST, 14793S), mouse anti-DYKDDDK-Tag (CST, 8146), rabbit anti-HA (CST, 3724), mouse anti-HA (CST, 2367), rabbit anti-His (CST, 12698), rabbit anti-Myc (CST, 13987), and mouse anti-Myc (CST, 2276); rabbit anti-Rig-I (CST, 3743), rabbit anti-MDA5 (CST, 5321), rabbit anti-MAVS (CST, 24930), rabbit anti-TRAF3 (CST, 33640), rabbit anti-IKKε (CST, 96794), rabbit anti-p-IKKε (CST, 8766), rabbit anti-TBK1 (CST, 3504), rabbit anti-p-TBK1 (CST, 5483), rabbit anti-IRF3 (CST, 4302), rabbit anti-p-IRF3 (CST, 29047), rabbit anti-IRF7 (CST, 72073), rabbit anti-p-IRF7 (CST, 24129), rabbit anti-LC3A/B (CST, 12741), rabbit anti-LC3B (CST, 2775), rabbit anti-SQSTM1/p62 (CST, 23214), rabbit anti-NDP52 (CST, 60732), rabbit anti-NIX (CST, 12396), mouse anti-PARKIN (CST, 4211), rabbit anti-PINK1 (CST, 6946), rabbit anti-BNIP3 (CST, 44060), rabbit anti-ATG5 (CST, 12994), rabbit anti-ATG7 (CST, 8558), rabbit anti-ATG12 (CST, 4180), and rabbit anti-ATG16L1 (CST, 8089) antibodies. Mouse anti-HSP60 (Proteintech, 2F1E7), rabbit anti-TOM20 (Proteintech, 11802-1-AP), rabbit anti-VDAC1 (Proteintech, 10866-1-AP), rabbit anti-GAPDH (Proteintech, 10494-1-AP), rabbit anti-β-tubulin (Proteintech, 10094-1-AP), rabbit anti-GST (Proteintech, HRP-66001), and rabbit anti-Lamin B1 (Proteintech, 12987-1-AP) antibodies were obtained from Proteintech (China). Anti-rabbit IgG HampL (Alexa Fluor 488; Invitrogen, A11034), anti-mouse IgG HampL (Alexa Fluor 555; Invitrogen, A30677), and anti-mouse IgG HampL (Alexa Fluor 594; Invitrogen, A21125) antibodies were purchased from Invitrogen. CQ phosphate (MCE, HY-17589), Z-VAD-FMK (MCE, HY-16658B), MG-132 (MCE, HY-13259), 3-MA (MCE, HY-19312), and CCCP (MCE, HY-100941) were acquired from MedChemExpress. NH4Cl (Sigma, 09718) and DMSO (Sigma, C6164) were purchased from Sigma‒Aldrich. SYBR Green (Thermo Scientific, 4385612) and NP-40 (Thermo Fisher Scientific, FNN0021) were obtained from Thermo Fisher Scientific. The dual-luciferase reporter assay system (Promega, E1910), nuclear extraction kit (Invent, SC-003), and IFN-β ELISA kit (Thermo Fisher Scientific, ES8RB) were purchased from the indicated companies.

### Plasmid construction

RIG-I, MDA5, MAVS, TRAF3, IKKε, TBK1, IRF3, IRF7 and ISG15 were amplified from DNA extracted from the tissues of pigs and inserted into the pcDNA-3.1 vector with an N-terminal HA tag. OPTN, NBR1, TOLLIP, NDP52 and SQSTM1/p62 were inserted into the pcDNA-3.1 vector, with an N-terminal Myc tag amplified from the *Homo sapiens* DNA sequence. Optimization of MGF_360-4L codons was suitable for humanized cell expression by Tsingke (Beijing, China) and subsequently cloned and inserted into pcDNA-3.1-HA, pcDNA-3.1-Flag and pcDNA-3.1-Myc vectors. Site-directed mutagenesis of pFlag-MGF_360-4L was performed via PCR to create ten different lysine-to-arginine mutations (201R, 208R, 209R, 257R, 290R, 295R, 320R, 321R, 327R, 329R) in pFlag-MGF_360-4L. Plasmids containing the IFN-β, ISRE and NF-κB promoters and TK plasmids along with pFlag-OAS1 were used for plasmid construction, as described previously (37). The primers used for plasmid construction are listed in **Table S1**.

### Generation of an ASFV strain lacking MGF_360-4L

To generate an ASFV strain lacking MGF_360-4L, ASFV CN/SC/2019 was used as the parental strain. Homologous arms flanking the MGF_360-4L gene were constructed, cloned, and inserted into a PUC19-based homologous recombination vector, which also contained an mCherry reporter gene under the control of the ASFV p72 promoter and an SV40 poly(A) signal. This vector was then transfected into PAMs, followed by ASFV CN/SC/2019 to achieve homologous recombination. The resulting recombinant virus, ASFV-△MGF_360-4L, was isolated through multiple passages and limited dilution, with mCherry fluorescence tracking used for selection. The deletion of MGF_360-4L and the identification of the recombinant virus were confirmed via PCR. The PCR primers used for constructing the transfer vector are listed in **Table S1**.

### One-step viral growth kinetics and HAD_50_ determination

PAMs were infected with ASFV CN/SC/2019 or ASFV-△MGF_360-4L at a MOI of 5. The virus was harvested from the infected cells at 1, 12, 24, and 48 hpi. The viral titer at each time point was determined via the 50% hemadsorption dose (HAD_50_) assay to calculate the virus growth curve. PAMs were cultured in a 96-well plate and infected with 10-fold serially diluted ASFV CN/SC/2019 or ASFV-△MGF_360-4L, with 8 replicates per dilution. Subsequently, 20 μL of 1% fresh red blood cells from pigs prepared in saline buffer were added to each well. The cells were incubated at 37°C and observed for hemadsorption (HAD) around infected cells within 7 dpi. The HAD_50_ was calculated via the Reed and Muench method (61).

### Dual-luciferase reporter assay

HEK293T cells were seeded into 24-well plates and transfected with plasmids containing ISRE, IFN-β NF-κB promoter elements, or pFlag-MGF_360-4L plasmids using jetPRIME (Polyplus, 101000046) for 12 hours. Concurrently, 0.01 μg of pRL-TK (Renilla luciferase) control reporter plasmid was included in the transfection system. Promoter activity was measured using the Promega Dual-Luciferase Reporter Assay System (Promega, E1910) according to the manufacturer’s instructions. Relative luciferase activity was calculated as the ratio of firefly luciferase activity to Renilla luciferase activity.

### Nuclear fractionation assay

HEK293T or HeLa cells were transfected with the pFlag-MGF_360-4L plasmid or control vector (VEC) and incubated for 20 hours. The cells were then stimulated with VSV for 6 hours. The cells were then collected and washed twice with ice-cold PBS. A specified amount of cytoplasmic extraction buffer was subsequently added, followed by vortexing in an ice bath for 5 minutes. The cytoplasmic extracts were collected from the supernatant. The pellet was resuspended in nuclear extraction buffer and separated using a centrifuge tube with a filter. The resulting fractions were subjected to Western blotting to verify the effect of nuclear-cytoplasmic separation. Lamin B1 and GAPDH were used as markers for the nuclear and cytoplasmic fractions, respectively.

### Transmission electron microscopy observation

HEK293T cells were transfected with either the pFlag-MGF_360-4L plasmid or empty vector and incubated for 24 hours. CCCP was added as a positive control. The cells were then harvested using 0.25% trypsin, centrifuged at 1,000 rpm for 10 minutes, fixed with 2.5% glutaraldehyde (Sigma, G6257) at a volume of 5 times that of the cell pellet, and incubated at 4°C for 12 hours. After fixation, the cells were sectioned and placed on metal copper grids, followed by staining with 2% osmium tetroxide and lead citrate. The samples were observed under a transmission electron microscope (Hitachi, TEM-HT7700).

### Indirect immunofluorescence assay

3D4/21 (CRL-2843) cells were seeded in 24-well plates containing coverslips (Wuhan, WHB-24-CS-10) and transfected with the specified plasmid for 24 hours. The cells were then washed twice with prechilled PBS, fixed with 4% paraformaldehyde solution (BBI, E672002-0500) at room temperature for 30 minutes, and subsequently incubated with 4% bovine serum albumin (BSA, Sigma Aldrich, A9647) at room temperature for 1 hour. The cells were permeabilized with 0.5% X-Triton 100 (Sigma‒Aldrich, T8787) for 10 minutes, followed by three washes with prechilled PBS. The cells were then stained with primary antibodies diluted to the specified concentration overnight and with goat anti-rabbit IgG H&L Alexa Fluor 488, goat anti-mouse IgG H&L Alexa Fluor 594, or goat anti-rabbit IgG H&L Alexa Fluor 555 secondary antibodies in the dark for 1 hour. After three additional washes with prechilled PBS, the coverslips were removed, and the cells were mounted with ProLong™ Glass Antifade Mountant (containing NucBlue™ stain) to visualize the nuclei. Colocalization was observed via a confocal fluorescence microscope (Zeiss).

### Co-IP and Western blot analysis

Whole-cell extracts were obtained by lysing cells in RIPA (Invitrogen, 89901) or NP-40 lysis buffer (20 mM Tris-HCl (pH 7.5), 150 mM NaCl, 1% NP-40, 1 mM EDTA, and protease inhibitor cocktail) containing 1 mM phenylmethylsulfonyl fluoride (PMSF) for 30 min at 4°C and centrifuging the lysate at 12,000 × g for 10 min to remove cell debris. For the IP reaction, the lysate was incubated with 2 µg of the indicated antibody or control IgG overnight at 4°C on a roller mixer. Protein A/G magnetic beads (Thermo, 88802) were subsequently added to capture the antibody-bound proteins. The protein A/G magnetic beads were washed with lysis buffer to remove nonspecific proteins, and then, the interacting proteins were detected via Western blot analysis. For Western blot analysis, the supernatants were then mixed with loading buffer and denatured at 100°C for 10 minutes. Proteins were separated by SDS‒ PAGE and transferred onto an nitrocellulose membrane. The membrane was blocked with 5% nonfat dry milk for 1 hour and then incubated with the specified primary antibody. Following washes with PBST, the membrane was incubated with the corresponding secondary antibody for 1 hour. The protein bands were visualized via a chemiluminescent substrate (Thermo Fisher Scientific, 32106).

### GST pull-down assay

The GST, GST-MGF_360-4L and His-tagged MDA5, p62, and OAS1 plasmids were transfected into BL21 competent cells following the manufacturer’s instructions. Protein expression was induced with IPTG at a final concentration of 1 mM. The GST-tagged proteins were purified using a GST Fusion Protein Purification Kit (GenScript, L00207), whereas the His-tagged proteins were purified using Ni-NTA affinity chromatography media (GenScript, L00250). The specified amounts of GST or GST-MGF_360-4L were incubated with glutathione resin at room temperature for 3 hours. After washing five times with ice-cold PBS, purified His-MDA5, His-p62, and His-OAS1 proteins were added and incubated at 4°C for 12 hours. Proteins were then eluted with elution buffer and subjected to Western blotting for analysis.

### RNA interference

To knockdown specific genes, small interfering RNAs (siRNAs) targeting the predicted target gene sequences were synthesized by Gemma Gene in Shanghai. HEK293T cells were cultured in 12-well plates and transfected with the corresponding siRNAs using jetPRIME transfection reagent, and the knockdown efficiency of the targets was then validated through Western blotting.

### RT‒qPCR

Total RNA from cells was extracted via the TRIzol method (Thermo Fisher Scientific, 15596018) according to the manufacturer’s instructions. Then, genomic DNA contamination was removed from the RNA template using a gDNA digester (MCE, HY-K0511A) following the manufacturer’s protocol, and reverse transcription was performed to obtain cDNA. The RT‒qPCR experiment was conducted with SYBR Green (Bio-Rad, 1725124) according to the manufacturer’s instructions. GAPDH was used as an internal control to normalize and quantify the expression of the target genes. The relative expression of each target gene was calculated via the 2^^−ΔΔCt^ method and normalized to the expression of GAPDH. Three independent experiments were performed.

### ELISA assay

To measure the p72 antibody levels in the isolated serum samples, ELISA was conducted following the manufacturer’s instructions (Ruiliwei, ASF.K001/5). Briefly, the serum samples were diluted 1:1 with sample diluent and incubated at 37°C for 1 hour, followed by washing with buffer four times. The enzyme-labeled antibody was added for binding and incubated at 37°C for 30 minutes. The TMB substrate was subsequently added and incubated in the dark for 15 minutes, followed by the addition of stop solution for the absorbance at 450 nm to calculate the sample blocking rate. The formula for S/N% is as follows: S/N% = (OD450 negative control – OD450 sample) / (OD450 negative control – OD450 positive control) × 100%, where S/N% ≤ 40% indicates negative, S/N% ≥ 50% indicates positive, and values outside of these two thresholds are considered unreliable. To measure IFN-β levels in the isolated serum samples, the samples were incubated at 4°C for 2.5 hours following the manufacturer’s instructions. After washing four times, the biotin-conjugated antibody was added, and the samples were incubated at room temperature for 1 hour. After a subsequent wash, streptavidin-HRP was added, and the mixture was incubated at room temperature for 45 minutes. TMB was then added, and the mixture was incubated for 30 minutes. Then, the reaction was terminated and the OD values at 450 nm were measured. The results are calculated using a standard curve derived from the equation y = 1766.9x + 0.124.

### Statistical analysis

Statistical analyses were conducted with Prism 9.3 software (GraphPad, San Diego, CA). The significance of differences between the experimental and control groups was analyzed by one-way analysis of variance (ANOVA). Two-tailed *p* values were calculated, and *p* values < 0.05 were considered to indicate statistical significance (* *p* < 0.05; ** *p* < 0.01, *** *p*< 0.001, and **** *p*< 0.0001). ns: nonsignificant difference.

### Data availability statement

All data generated or analyzed during the current study are included in this article and its supplementary information files.

## Acknowledgments

This research was financially supported by the National Natural Science Foundation of China (Grant No. 32072830), the National Key Research and Development Program (Grant No. 2021YFD1800101), the Gansu Provincial Major Project for Science and Technology Development (Grant No. 22ZD6NA001), the Science Fund for Creative Research Groups (22JR5RA024) and Special Project (22CX8NA011) of Gansu Province, the Central Public-interest Scientific Institution Basal Research Fund (Grant No. CAAS-ZDRW202409) and the Innovation Program of the Chinese Academy of Agricultural Sciences (CAAS-ASTIP-2021-LVRI).

